# Systematic comparison of genetically encoded F-actin probes for visualizing actomyosin dynamics *in vivo* using *C. elegans*

**DOI:** 10.64898/2026.11.03.763946

**Authors:** Taylor N. Medwig-Kinney, Pu Zhang, Amy Liu, Michael E. Werner, Naveen K. Chana, Amy Shaub Maddox, Bob Goldstein

**Affiliations:** Department of Biology, University of North Carolina at Chapel Hill, Chapel Hill, NC, USA; Department of Biology, University of Wisconsin–La Crosse, La Crosse, WI, USA; Center for Life Sciences, Peking University, Beijing, China; Lineberger Comprehensive Cancer Center, University of North Carolina at Chapel Hill, Chapel Hill, NC, USA

**Author notes:** **These authors contributed equally.** (Taylor N. Medwig-Kinney, Pu Zhang). **Address correspondence to:** Taylor N. Medwig-Kinney.

## Abstract

Genetically encoded F-actin probes enable visualization of actin organization and dynamics in living cells, but their performance and effects on cellular function can vary by probe, fluorescent protein, and biological context. Here, we systematically compared three F-actin probes – F-tractin, LifeAct, and Utrophin – fused to mScarlet-I, mNeonGreen, or mStayGold *in vivo*, using *Caenorhabditis elegans*. We used single-copy transgenic strains and evaluated fluorescence intensity, photostability, developmental and reproductive outcomes, cytokinetic ring closure, and probe localization across diverse cellular contexts. mStayGold provided the greatest brightness and photostability, making it well suited for long-term time-lapse imaging, whereas mNeonGreen offers an alternative for applications requiring photobleaching. Most of the strains exhibited minimal detectable effects on development and cellular function and outperformed previously generated strains tested. Probe performance varied across tissues and developmental stages. F-tractin generally exhibited lower fluorescence intensity than LifeAct and Utrophin, particularly in embryos. Although the probes largely exhibited subcellular localization patterns consistent with phalloidin staining, differences in their labeling of specific F-actin structures were observed. Together, these results establish a practical framework for selecting genetically encoded F-actin probes for *C. elegans*, they suggest best candidates for other *in vivo* systems, and they highlight the importance of considering both fluorescent protein and actin-binding probe when designing live-cell imaging experiments.

**Significance Statement:**

- This collection of genetically encoded F-actin probe strains in *C. elegans*, together with modular plasmids for generating customized probes, provides accessible resources for a broad range of imaging applications.
- All tested probes support improved retention of normal biological function, and vary in fluorescence intensity, photostability, and the labeling of some actin structures, revealing trade-offs that inform probe selection.
- Evaluating probes across developmental stages and cellular processes demonstrates that the optimal probe depends on the biological context and experimental goals.

## Introduction

Actomyosin networks are fundamental to cell architecture and drive essential cellular processes throughout development, including cytokinesis, cell shape changes, cell migration, and tissue remodeling (see Pollard and Cooper, 2009, for review). Visualizing filamentous actin (F-actin) in living cells is essential for understanding how actin dynamics contribute to these processes. Fluorescently labeled phalloidin is widely considered a gold-standard tool for visualizing F-actin, but it disrupts F-actin dynamics and its use typically requires fixation and permeabilization, limiting its use to a static snapshot of actin organization rather than allowing actin dynamics to be monitored in living cells over time (Wulf *et al*., 1979; Cooper, 1987). Directly tagging actin with GFP disrupts normal dynamics, so F-actin binding proteins or fragments of them are usually used (see Melak *et al*., 2017, for review). Genetically encoded F-actin probes enable researchers to visualize actin structures and dynamics *in vivo*, and their performance can vary, likely based on how each probe interacts with F-actin, the fluorescent protein to which it is fused, and the biological context in which it is expressed.

Although F-actin markers have been systematically compared in cultured cells (Belin *et al*., 2013, 2014), such controlled comparisons in animal systems *in vivo* remain limited. The effects of markers on normal development and cellular function must be considered when interpreting their localization and dynamics *in vivo*. In *Drosophila* oogenesis, multiple probes have been compared systematically, although F-actin organization and/or animal fertility at commonly used temperatures were disrupted under the conditions and expression levels used (Spracklen *et al*., 2014). Furthermore, the fluorescent proteins and linker sequences used in genetically encoded probes can influence their subcellular localization and the F-actin populations they label (Lemieux *et al*., 2014). In addition, LifeAct and Utrophin-derived probes have been shown to induce abnormal actin assemblies and interfere with actin dynamics, with effects that vary depending on the probe and cellular context (Du *et al*., 2015). These findings highlight the need to systematically evaluate probe performance, considering a broad spectrum of potential effects on biological function.

Here, we generated strains with various F-actin probes and fluorescent protein tags, and we compared fluorescence intensity, photostability, effects on development and cellular function, as well as performance across diverse F-actin-dependent cellular processes. Our results reveal trade-offs among probe properties, demonstrate that the optimal probe depends on the biological context and experimental goals, and provide practical guidance and adaptable resources for F-actin imaging.

## Results

### Design of tagged probes and strains for systematic comparisons

We generated a collection of *C. elegans* strains expressing three commonly used, genetically encoded F-actin probes: F-tractin, LifeAct, and the CH domains of Utrophin (hereafter referred to as Utrophin for brevity). F-tractin (43 amino acids from rat ITPKA) and LifeAct (17 amino acids from budding yeast Abp140) are both short peptides that bind to overlapping regions of F-actin at the interface between adjacent actin subunits (Riedl *et al*., 2008; Johnson and Schell, 2009; Belyy *et al*., 2020; Shatskiy *et al*., 2025). In contrast, Utrophin uses a larger actin-binding domain comprising two calponin homology (CH) domains (Burkel *et al*., 2007). Each of these probes was fused to one of three fluorescent proteins: mNeonGreen, mStayGold, or mScarlet-I (**Figure 1A**). These fluorescent proteins possess distinct spectral properties and photostability characteristics, enabling probe selection for different imaging applications, including multicolor imaging. To facilitate direct comparisons while changing just one variable at a time, we used a common expression strategy for all probe variants. We optimized coding sequences for expression in the germline via a published germline optimization method developed by Dan Dickinson (Fielmich *et al*., 2018), removed piRNA homology (Priyadarshini *et al*., 2022), and placed each probe under the control of the *mex-5* promoter to drive strong expression in the germline and early embryos.

**Figure 1.**
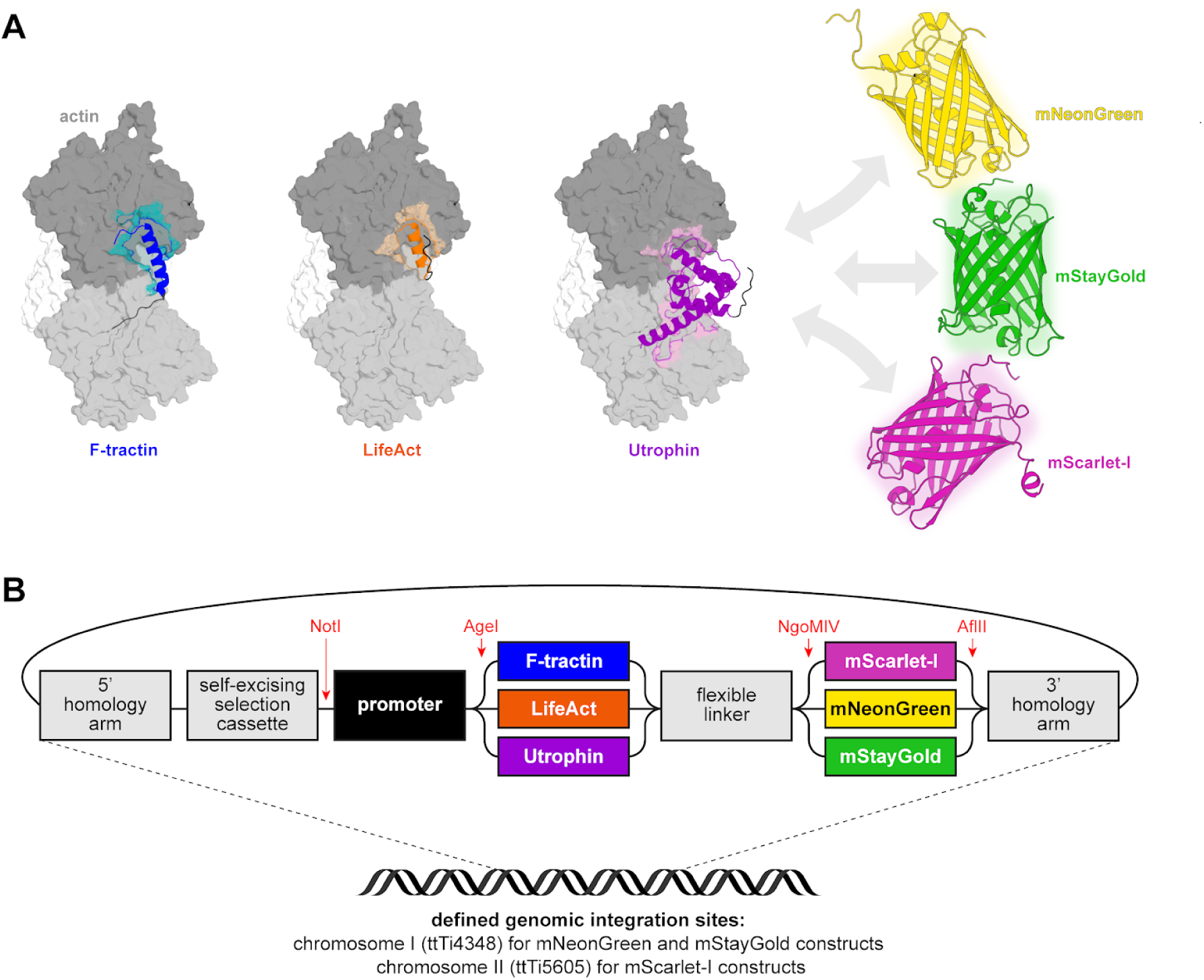
Design of modular, single-copy genetically encoded F-actin probes. (A) Protein structures of actin, the F-actin probes F-tractin, LifeAct, and Utrophin, and the fluorescent proteins mNeonGreen, mStayGold, and mScarlet-I. (B) Schematic of the CRISPR plasmids designed for single-copy integration of fluorescently tagged F-actin probes. Each construct contains 5’ and 3’ homology arms flanking a self-excising selection cassette and a probe expression module consisting of a promoter, an F-actin probe (F-tractin, LifeAct, or Utrophin), a flexible linker, and a fluorescent protein (mNeonGreen, mStayGold, or mScarlet-I). Coding sequences were optimized for germline expression. Constructs containing mNeonGreen or mStayGold target the defined genomic integration locus ttTi4348 on chromosome I. Constructs containing mScarlet-I target ttTi5605 on chromosome II. Unique restriction sites (shown in red) flanking the promoter and fluorescent protein modules enable modular exchange of these components.

To generate a standardized collection of probe-expressing strains, we employed plasmid-based CRISPR/Cas9-mediated genome engineering, using the self-excising cassette selection strategy (Dickinson *et al*., 2015). Single-copy transgenes were integrated at the MosSCI sites ttTi4348 on chromosome I (for the mNeonGreen and mStayGold plasmids) and ttTi5605 on chromosome II (for the mScarlet-I plasmids) to minimize disruption of endogenous gene function and reduce variability associated with transgene copy number and random integration (Frøkjær-Jensen *et al*., 2008, 2012). We designed these repair templates to be modular through the inclusion of unique restriction sites that facilitate the exchange of promoter and fluorescent protein modules (**Figure 1B**). This design allows adaptation of the constructs for different expression patterns and imaging requirements.

### Fluorescence intensity and photostability vary among genetically encoded F-actin probes

Important considerations for selecting an F-actin probe for imaging are its fluorescence intensity and photostability. To maximize comparability across strains, we analyzed embryos at the four-cell stage, when all embryonic cells can be visualized in the same focal plane, as done previously when comparing fluorescent proteins (Heppert *et al*., 2016). We used morphology of one of the cells, EMS, to closely synchronize embryos to within 10 minutes in age. We compared our newly engineered strains with two commonly used strains: LifeAct::mKate2 driven by the *mex-5* promoter (Reymann *et al*., 2016) and LifeAct::GFP driven by the *pie-1* promoter (Redemann *et al*., 2010). For each fluorescence color channel, images were acquired using equivalent imaging conditions and displayed with equivalent settings (**Figure 2A-D**).

**Figure 2.**
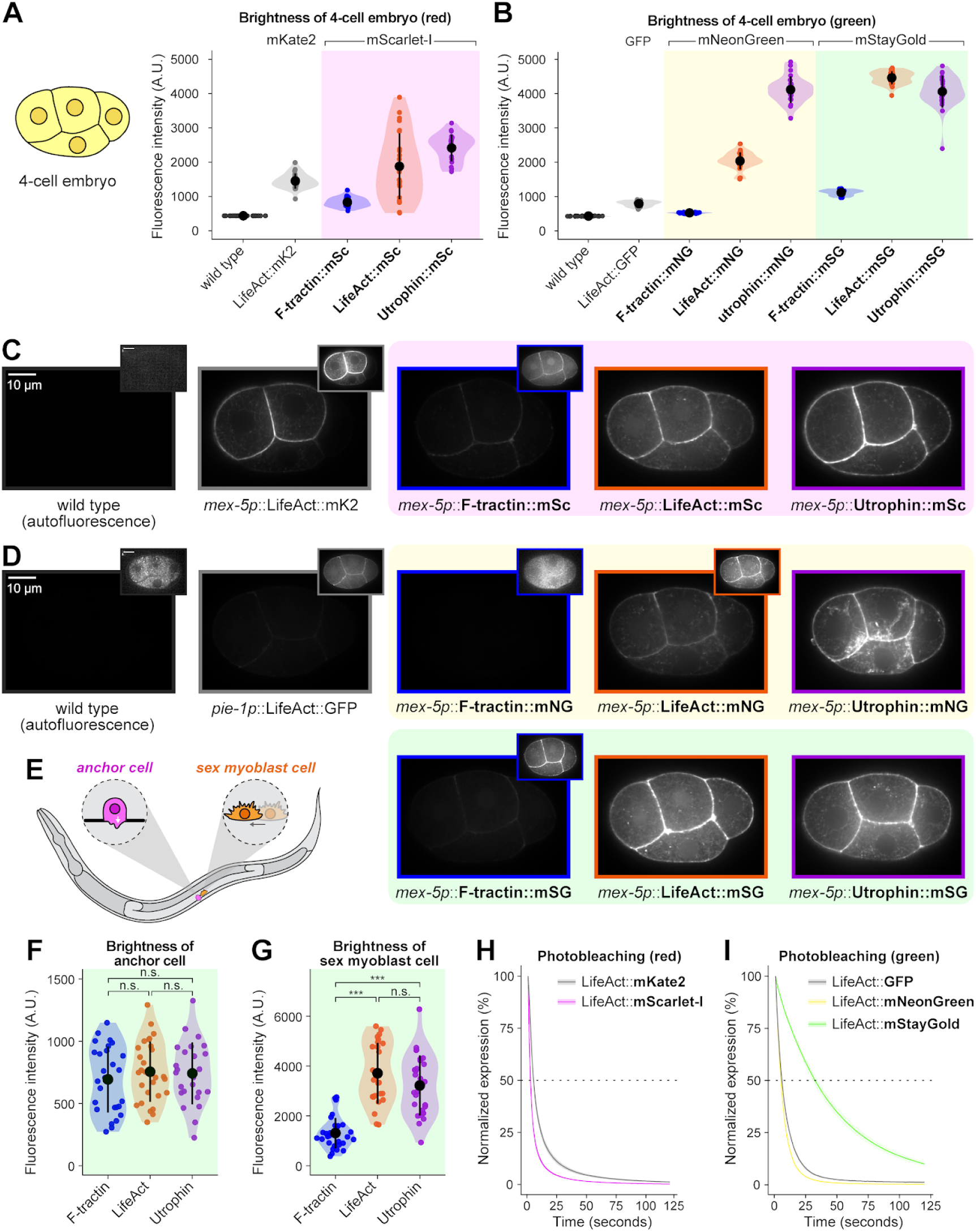
Comparison of fluorescence intensity and photostability of genetically encoded F-actin probes. (A, B) Quantification of fluorescence intensity (measured in arbitrary units) of four-cell stage embryos expressing nine newly generated F-actin probes (bolded), each expressed under the germline/embryo promoter *mex-5* and comprising one of three actin-binding probes (F-tractin, LifeAct, or Utrophin) tagged with one of three fluorescent proteins (mScarlet-I, mNeonGreen, or mStayGold). Wild type animals and previously established probes (*mex-5p*::LifeAct::mKate2 and *pie-1p*::LifeAct::GFP) are included for comparison. Whole-embryo fluorescence intensity was measured at a single optical plane through the embryo midpoint. (C-D) Representative micrographs of the indicated strains acquired under matched imaging conditions and displayed using identical brightness and contrast settings for each fluorescent channel – red (C) and green or yellow (D). Insets show images displayed at increased brightness to reveal dim fluorescence. Green-channel fluorescence in wild-type animals reflects autofluorescence under the same settings as used for GFP, mNeonGreen, and mStayGold. (E) Schematic illustrating the biological context of the anchor cell and sex myoblast. (F, G) Quantification of fluorescence intensity of mStayGold-tagged F-actin probes expressed under the *cdh-3* promoter in the anchor cell (F) and the *hlh-8* promoter in the sex myoblasts (G). (H, I) Photobleaching curves for LifeAct-tagged probes, comparing mKate2 and mScarlet-I (H) and GFP, mNeonGreen, and mStayGold (I). Statistical significance for panels F and G was determined using Welch’s one-way ANOVA followed by Games-Howell post-hoc tests. Statistical results for panels A, B, H, and I are presented in Supplemental Figure S1.

The previously established *mex-5p*::LifeAct::mKate2 strain exhibited moderate fluorescence intensity, and signal intensity varied among the four cells of each embryo. Fluorescence intensity was consistently higher in ABa and ABp cells, which are AB descendants, than in EMS and P_2_, which are P_1_ descendants. Our F-tractin::mScarlet-I strain was 43% dimmer than LifeAct::mKate2. In contrast, our LifeAct::mScarlet-I and Utrophin::mScarlet-I probes were significantly brighter than LifeAct::mKate2 (29% and 66% greater fluorescence intensities, respectively) (**Figure 2A,C; Supplemental Figure S1A**).

We next compared the mNeonGreen fluorescent probes with the existing *pie-1p*::LifeAct::GFP strain. All our new strains except F-tractin::mNeonGreen exhibited significantly greater fluorescence intensity than the GFP-tagged strain. LifeAct::mNeonGreen was more than 1.5-fold times as bright, whereas Utrophin::mNeonGreen was more than fourfold as bright. The F-tractin::mStayGold and LifeAct::mStayGold strains were approximately twice as bright as their respective mNeonGreen-tagged counterparts. Interestingly, however, this trend did not hold for Utrophin; the mNeonGreen- and mStayGold-tagged versions were similar in fluorescence intensity (**Figure 2B,D; Supplemental Figure S1A**).

Although all probe-coding sequences were optimized for germline expression, we considered that the lower fluorescence intensity of F-tractin compared with the other probes, regardless of fluorescent protein used, could reflect germline-specific silencing. We examined probe expression in two postembryonic somatic cell types that exhibit actin-dependent behaviors: the anchor cell, a specialized uterine cell that invades the basement membrane separating the uterus and vulval epithelia to create a passage for egg laying (Sherwood and Sternberg, 2003), and the sex myoblast, which migrates anteriorly toward the center of the body to generate vulval and uterine muscles required for egg laying (Sulston and Horvitz, 1977) (**Figure 2E**). We expressed the mStayGold-tagged probes in the anchor cell using *cis*-regulatory elements from the *cdh-3* promoter. No significant differences in fluorescence intensity were detected among the three probes (**Figure 2F**). We then expressed the same probes in sex myoblasts using the *hlh-8* promoter. In this context, F-tractin::mStayGold was approximately 60-65% dimmer than the LifeAct and Utrophin probes, whereas LifeAct and Utrophin did not differ significantly in fluorescence intensity (**Figure 2G**). These results suggest that the low fluorescence intensity of F-tractin is not explained solely by germline-specific silencing, and that its expression, stability, and/or fluorescence intensity is influenced by cell context.

Finally, we compared photobleaching rates among the LifeAct strains to evaluate the effects of fluorescent protein identity on probe photostability. Photobleaching was measured in four-cell-stage embryos. Among the red fluorescent probes, mScarlet-I photobleached slightly faster than mKate2 (**Figure 2H; Supplementary Figure S1B**). Among the yellow or green fluorescent probes, mNeonGreen photobleached slightly faster than GFP, whereas mStayGold was significantly more photostable, as expected (Ando *et al*., 2024) (**Figure 2I; Supplementary Figure S1B**).

### New F-actin probes minimally perturb development and cellular function

An important consideration when using genetically encoded probes is whether their expression perturbs the biological processes they are intended to visualize. Because F-actin is involved in diverse cellular functions, probes that alter actin organization or dynamics may introduce unintended effects on development and cell behavior. We therefore assessed the effects of our newly generated F-actin probes on organismal development and cellular processes, comparing them with existing probe strains and appropriate controls.

As an initial measure of organismal health, we quantified brood size over a 96-hour period. A fluorescently tagged membrane marker, *mex-5p*::mNG::PLCδ-PH (Heppert *et al*., 2016), was included as an additional control to assess developmental effects associated with expression of a fluorescently tagged protein that does not directly bind F-actin. Compared with wild-type (N2) animals, the membrane-associated PLCδ-PH control exhibited highly variable, and 24% reduced on average, brood size. The two previously established F-actin probe strains also exhibited reduced brood sizes: *mex-5p*::LifeAct::mKate2 produced an average of 37% fewer embryos, and *pie-1p*::LifeAct::GFP produced 33% fewer embryos. Among our newly generated F-actin probe strains, only the Utrophin::mScarlet-I strain exhibited a significant reduction in brood size (14%) relative to wild-type animals. The remaining strains showed no significant reduction, with some instead exhibiting a slight but statistically significant increase (**Figure 3A**). Overall, these results suggest that our new probes had limited effects on reproductive output.

**Figure 3.**
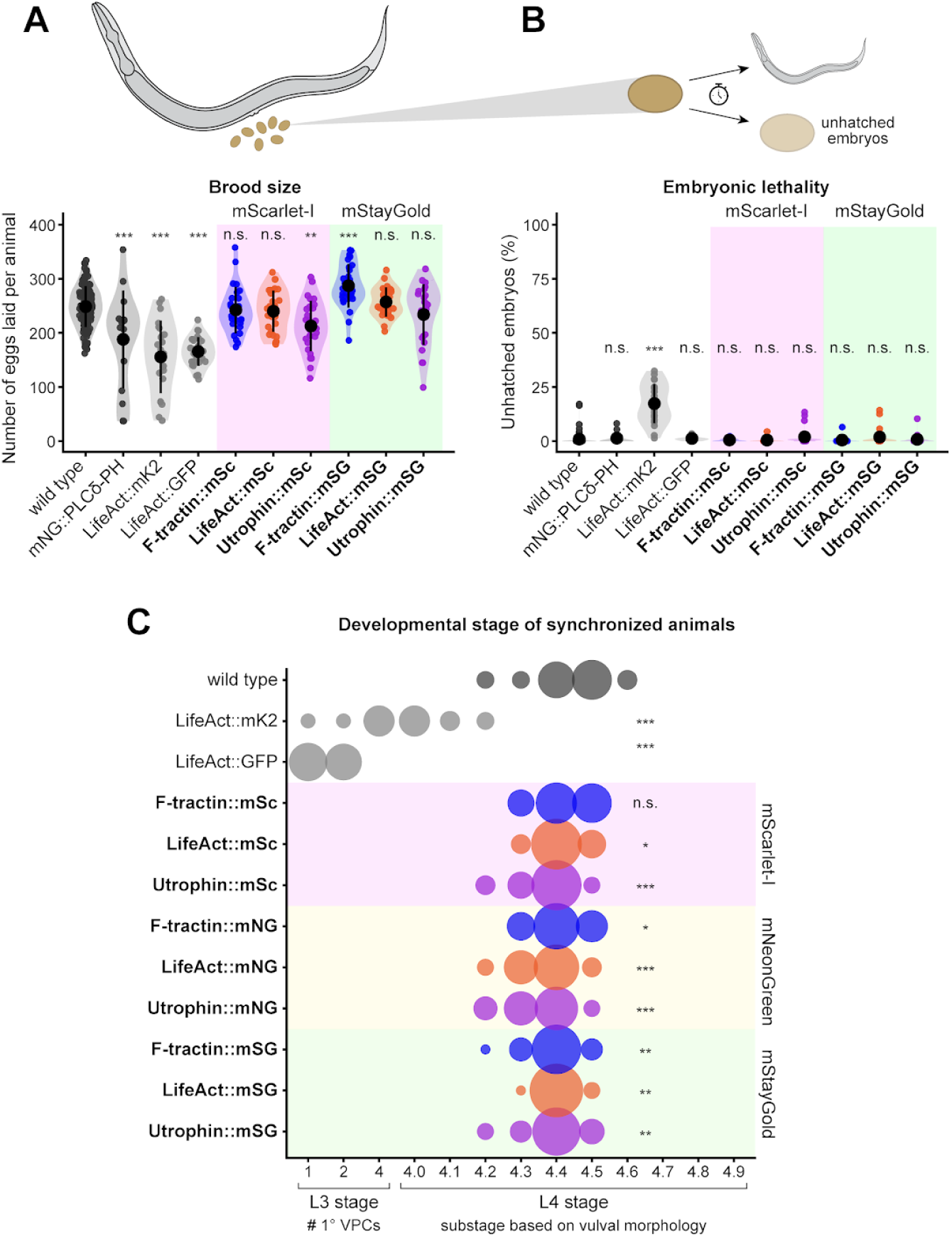
Effects of F-actin probe expression on fertility and development. (A) Brood size and (B) embryonic lethality of strains expressing F-actin probes, including wild-type and *mex-5p*::mNG::PLCδ-PH controls, the two previously established probe strains (*mex-5p*::LifeAct::mKate2 and *pie-1p*::LifeAct::GFP), and newly generated mScarlet-I- and mStayGold-tagged probe strains. (C) Bubble plot showing the distribution of L3 and L4 developmental substages, assessed by vulval development, in synchronized animals 39 h after release from L1 arrest at 20°C. L3 substages were classified by the number of primary vulval precursor cells (1° VPCs), whereas L4 substages were determined by vulval morphology using an established staging system. Bubble size represents the proportion of animals at each substage. Statistical significance was determined using one-way ANOVA followed by Dunnett’s post-hoc test (A, B) or Wilcoxon rank-sum tests with Holm correction for multiple comparisons (C), comparing each strain to wild type.

We next assessed embryonic viability by measuring the percentage of embryos that failed to hatch within 48 hours of being laid. Wild-type animals exhibited an average embryonic lethality rate of less than 1%. *mex-5p*::LifeAct::mKate2 exhibited significantly increased embryonic lethality, with approximately 17% of embryos remaining unhatched after 48 hours. In contrast, neither *pie-1p*::LifeAct::GFP nor any of our newly generated probes exhibited a significant increase in embryonic lethality relative to wild-type animals (**Figure 3B**). Together, these findings indicate that the increased embryonic lethality observed with *mex-5p*::LifeAct::mKate2 is not a general consequence of expressing a genetically encoded fluorescent F-actin probe.

We next examined whether probe expression affected postembryonic growth. To assess growth over time, we synchronized animals by collecting embryos laid within a two-hour window and measured their body lengths 48, 72, and 96 hours later. At 48 hours, all F-actin probe strains except F-tractin::mStayGold exhibited significantly smaller body sizes than the wild type control. The *mex-5p*::LifeAct::mKate2 strain reached only 71% of the mean wild-type body length. Among our newly generated strains, those expressing mScarlet-I-tagged probes reached 83-86% of the wild-type mean, whereas mStayGold-tagged strains reached 91-95%. These differences narrowed over time. By 96 hours, all newly generated probe strains were within 6% of the mean wild-type animal body length, whereas *mex-5p*::LifeAct::mKate2 animals remained approximately 10% smaller. Thus, although early growth was reduced across probe strains, the effects were less pronounced among our new probes and largely diminished by 96 hours (**Supplemental Figure S2**).

To further investigate developmental progression while distinguishing postembryonic delays from potential effects arising during embryogenesis, we performed a complementary experiment using animals synchronized by hypochlorite treatment (Porta-de-la-Riva *et al*., 2012). After collection, embryos were allowed to hatch and arrest at the L1 larval stage before being placed on food to resume development. To assess developmental progression more precisely, we used a staging system based on the morphology of the developing vulva that divides the L4 larval stage into ten substages (Mok *et al*., 2015). Animals were examined after 39 hours at 20°C, when wild-type animals had reached the L4.4 stage on average (**Figure 3C**). All newly generated F-actin probe strains reached an average vulval substage of 4.4, comparable to wild-type. In contrast, the previously established LifeAct strains exhibited delayed development. Animals expressing *mex-5p*::LifeAct::mKate2 averaged the earliest L4 substage (L4.0) and showed a broader distribution of developmental stages, including some animals that remained in the L3 stage. The *pie-1p*::LifeAct::GFP strain exhibited the most pronounced delay, with no animals having progressed to the L4 stage by the time of examination. Because L3 stage animals cannot be assigned the same vulval substages used to distinguish L4 development, we additionally assessed the division status of the primary-fated vulval precursor cell (1° VPC) lineage, which progresses through two divisions during the L3 stage (Schindler and Sherwood, 2013). Animals expressing *pie-1p*::LifeAct::GFP had only one or two cells in the 1° VPC lineage, consistent with development at the early-to-mid-L3 stage. Together, these results demonstrate that the previously established LifeAct strains exhibit pronounced delays in postembryonic development, whereas animals expressing the newly generated probes showed mean developmental stages within one substage of wild-type animals under the conditions tested.

To determine whether these developmental effects were accompanied by changes in cellular function, we next measured cytokinetic ring closure time during the first embryonic division, defined as the interval between initial cleavage furrow ingression and ring closure. Using the mNeonGreen::PLCδ-PH strain as a control, we found no significant differences in ring closure time in embryos expressing *pie-1p*::LifeAct::GFP or any of the newly generated mStayGold-tagged F-actin probes (**Figure 4A,C**). Among the mScarlet-I-tagged probes, F-tractin::mScarlet-I was too dim and photobleached too rapidly to permit reliable measurements. LifeAct::mScarlet-I expression did not significantly affect ring closure time, whereas Utrophin::mScarlet-I expression was associated with a small but statistically significant delay of approximately 13 seconds, or 7% slower on average, relative to the control (**Supplemental Figure S3A,B**). In contrast, embryos expressing *mex-5p*::LifeAct::mKate2 exhibited a pronounced delay in cytokinetic ring closure, averaging 108 seconds, or 54%, longer than the control (**Figure 4A,C**). In these experiments, we also observed an apparent strong asymmetry in the fluorescence distribution of *mex-5p*::LifeAct::mKate2 during the first embryonic division, consistent with the asymmetry observed for this marker at the four-cell stage (**Figure 2A,C**) but not seen in any of the new strains (**Figure 4B,C**). Collectively, these results demonstrate that the newly generated F-actin probes generally have modest to no effects on organismal development, reproductive function, and cytokinesis.

**Figure 4.**
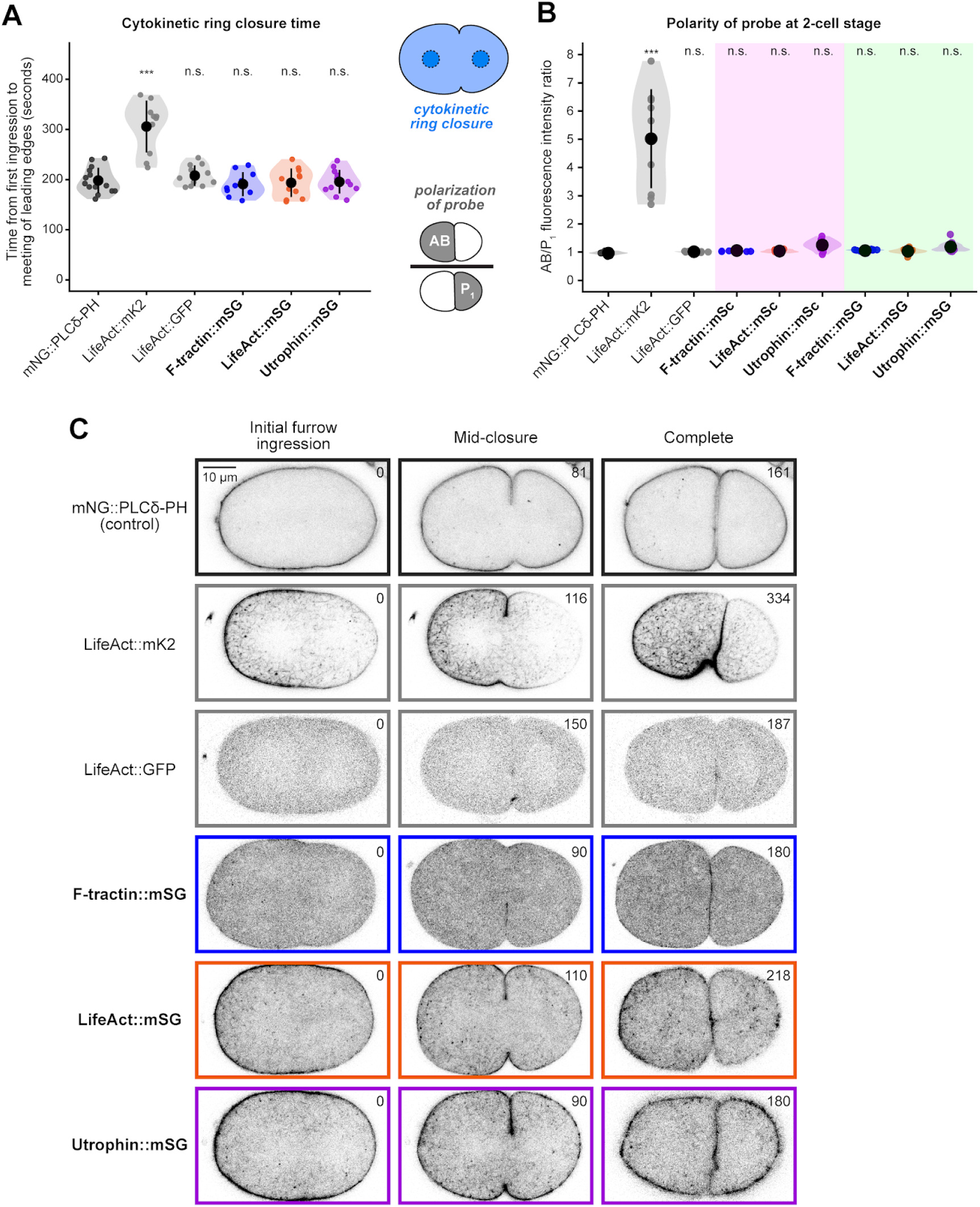
Effects of F-actin probes on cytokinetic ring closure and fluorescence polarity. (A) Quantification of cytokinetic ring closure time, measured in seconds from initial furrow ingression to the meeting of the leading edges, in embryos expressing mStayGold-tagged F-actin probes, the previously established probes *mex-5p*::LifeAct::mKate2 and *pie-1p*::LifeAct::GFP, and the *mex-5p*::mNG::PLCδ-PH control. (B) Quantification of fluorescence polarity at the two-cell stage, expressed as the ratio of fluorescence intensity of the AB cell to that in the P_1_ cell (AB/P_1_). Statistical significance was determined using one-way ANOVA followed by Dunnett’s post-hoc test, comparing each strain to the *mex-5p*::mNG::PLCδ-PH control. (C) Representative micrographs of embryos expressing the indicated probes at initial furrow ingression, mid-closure, and completion of cytokinetic ring closure. Fluorescence signals are displayed using an inverted lookup table. Time in seconds is indicated in each image.

### Genetically encoded F-actin probes enable visualization of actin dynamics in diverse cellular contexts

To determine how well our markers matched F-actin distribution recognized by fluorescent phalloidin, we compared the subcellular localization of the newly generated mStayGold-tagged F-actin probes with Alexa Fluor 488-conjugated phalloidin staining and the two previously established LifeAct-expressing strains in four-cell-stage embryos (**Figure 5**). We examined probe localization at cell-cell contacts, along exposed cell cortices, and within the cytoplasm, including perinuclear structures. Overall, with the exception of *mex-5p*::LifeAct::mKate2, which exhibited substantially uneven fluorescence intensity between different embryonic cells, the genetically encoded probes exhibited localization patterns that were largely consistent with phalloidin staining. Perinuclear puncta and cortical F-actin bundles were observed in embryos expressing all three mStayGold-tagged probes, although these structures were more readily distinguishable in LifeAct::mStayGold- and Utrophin::mStayGold-expressing embryos than in F-tractin::mStayGold-expressing embryos, likely reflecting their higher fluorescence intensity and greater contrast against background autofluorescence, which is elevated at 488 nm illumination compared to other wavelengths (Heppert *et al*., 2016). Additionally, Utrophin::mStayGold exhibited noticeably reduced nuclear fluorescence compared with the other probes.

**Figure 5.**
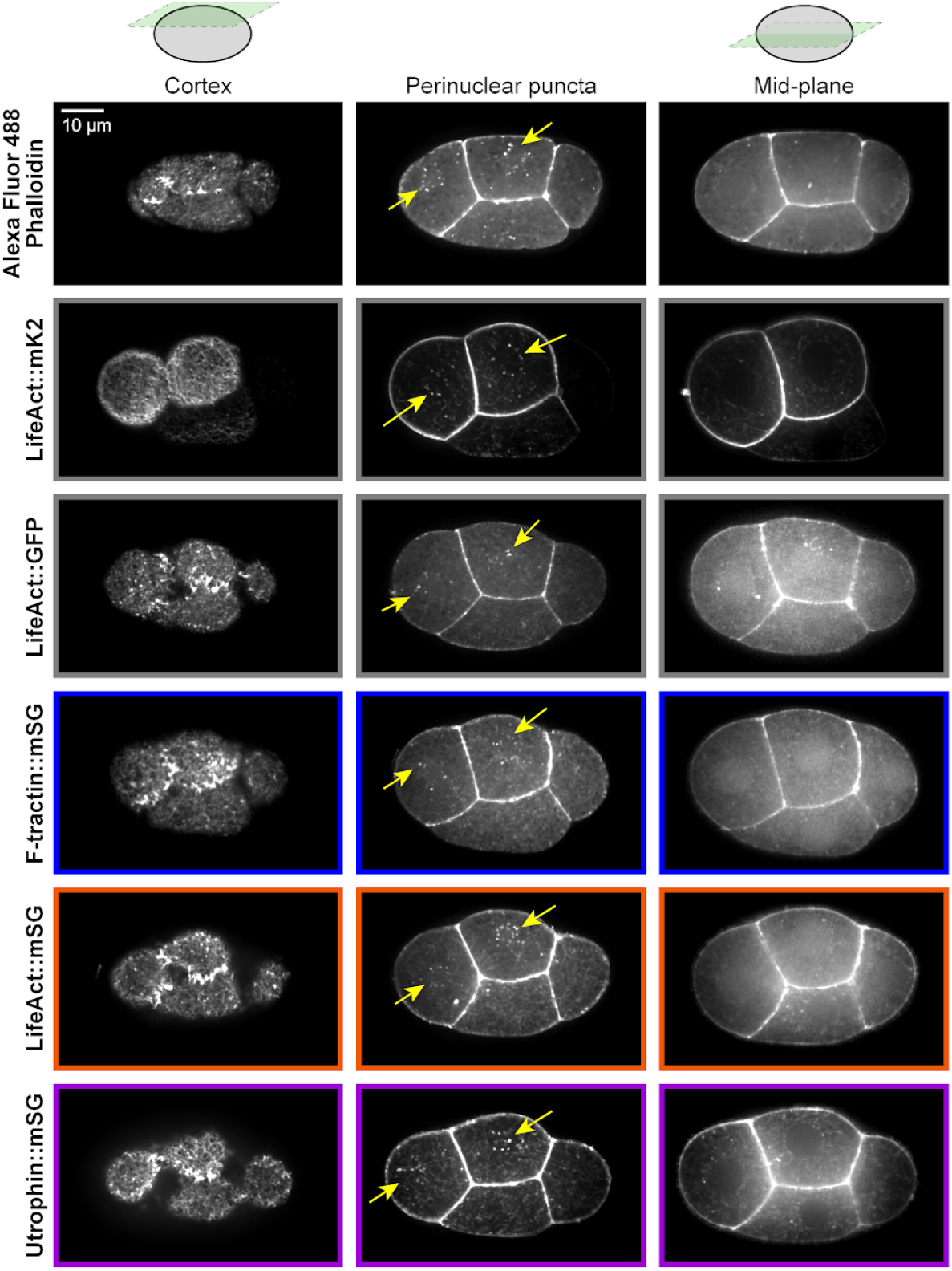
Comparison of subcellular localization of genetically encoded F-actin probes in four-cell stage embryos. Subcellular localization of newly generated mStayGold-tagged F-actin probes, with Alexa Fluor 488 phalloidin staining and previously established probes (*mex-5p*::LifeAct::mKate2 and *pie-1p*::LifeAct::GFP) included for comparison. Representative images of the cortical plane, perinuclear plane, and mid-plane through the nuclei.

To assess the utility of these strains for visualizing F-actin dynamics across different cellular behaviors and contexts, we examined F-actin network dynamics in multiple cells that perform a diverse set of motile or contractile F-actin-dependent behaviors, and whether the probes compromised these behaviors. We first examined apical constriction during *C. elegans* gastrulation using the embryonically expressed mStayGold-tagged probes (**Figure 6A**). Apical constriction involves contraction of the apical cortex, reducing the apical surface area and driving changes in cell shape (Martin and Goldstein, 2014). We focused on the endodermal precursor cells Ea and Ep, which internalize during gastrulation, a process involving dynamic reorganization of the cortical actin cytoskeleton (Goldstein and Nance, 2020; Zhang *et al*., 2023). To assess whether probe expression affected internalization, we measured its timing relative to division of the neighboring MSx cell using differential interference contrast (DIC) microscopy. Internalization occurred approximately 16 minutes after MSx division on average, with no significant differences between embryos expressing the mStayGold-tagged probes and wild-type controls (**Figure 6B**).

**Figure 6.**
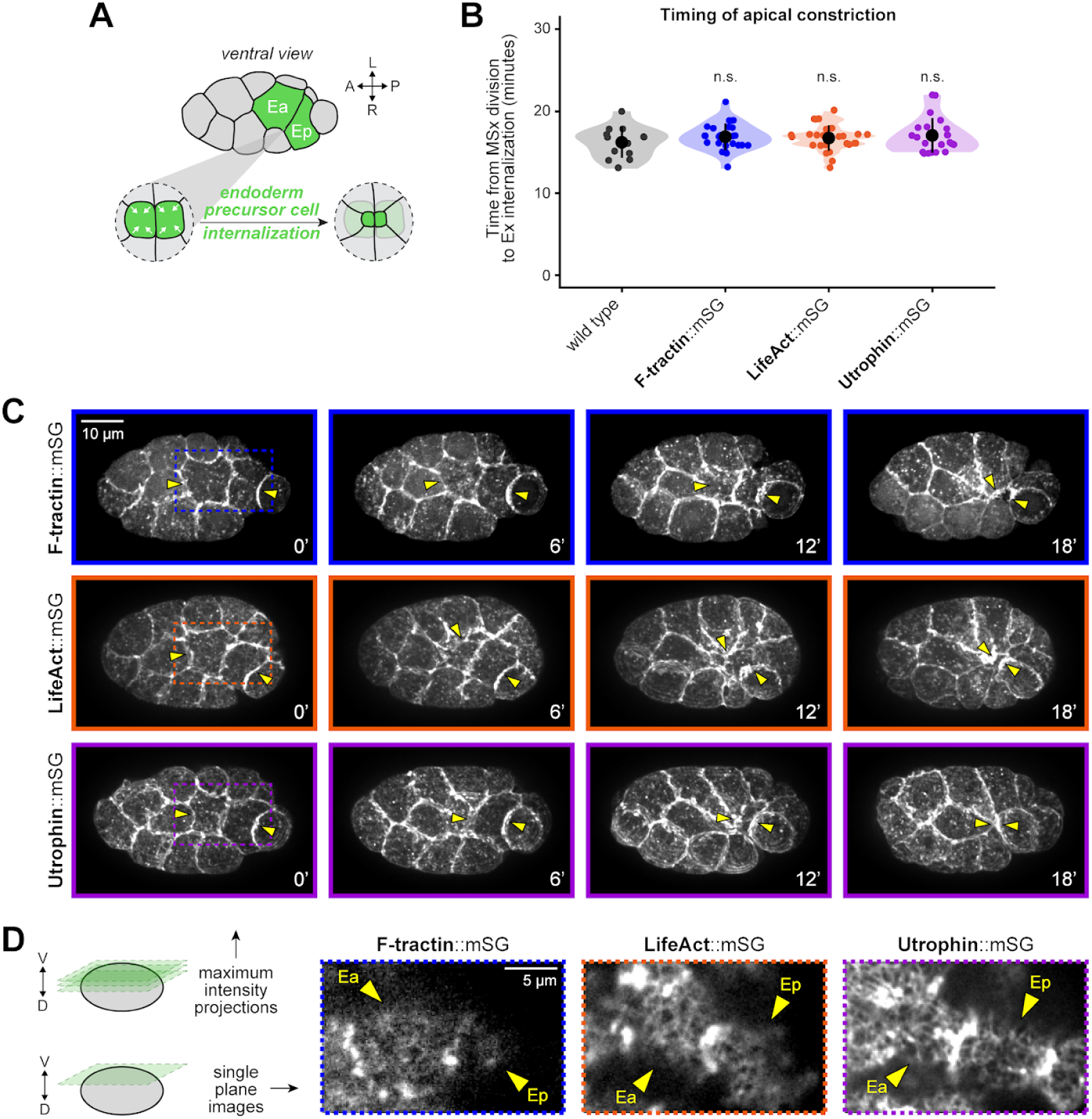
F-actin probes during apical constriction of endoderm precursor cells. (A) Schematic depicting apical constriction and internalization of the endoderm precursor cells Ea and Ep (highlighted in green), shown from a ventral view. (B) Timing of Ea and Ep internalization relative to division of the MS descendants (MSx) in embryos expressing the indicated mStayGold-tagged F-actin probes. Statistical significance was determined using one-way ANOVA followed by Dunnett’s post-hoc test, comparing each strain to wild type. (C) Representative shallow maximum intensity projections of embryos at 0, 6, 12, and 18 minutes after MSx cell division. Yellow arrowheads indicate Ea and Ep, and dashed boxes indicate the regions shown at higher magnification in (D), which displays a single z-plane.

We next imaged F-actin dynamics during apical constriction to evaluate the utility of these probes for time-lapse imaging through multiple focal planes and time points. The mStayGold-tagged probes were best suited for this application because of their brightness and photostability (**Figure 6C**). We also acquired time-lapse sequences using the mScarlet-I-tagged LifeAct and Utrophin strains (**Supplemental Figure S4A**); however, rapid photobleaching limited their utility for this degree of extended, multi-plane imaging, and the F-tractin::mScarlet-I strain in particular was too dim and photobleached too rapidly to obtain usable time-lapse sequences. We additionally imaged the mNeonGreen-tagged probes at mid-apical constriction (**Supplemental Figure S4B**). Among the mStayGold-tagged probes, F-tractin fluorescence was the dimmest, with fine F-actin structures difficult to resolve, whereas LifeAct and Utrophin provided better imaging of these structures. Utrophin labeling also appeared to reveal more clearly visible F-actin network patterns at the apical cortex than LifeAct (**Figure 6D**)

We next examined F-actin organization in the anchor cell as it invades the basement membranes separating the uterus and vulval epithelium (Sherwood and Sternberg, 2003) (**Figure 7A**). During this process, the anchor cell produces F-actin-rich invasive protrusions at its basal surface, where they promote localized basement membrane breaching (Hagedorn *et al*., 2013). To visualize these dynamics specifically within the anchor cell, we generated three additional strains expressing the F-actin probes under the control of the anchor cell-specific *cis*-regulatory elements from the *cdh-3* promoter and minimal *pes-10* promoter. To determine whether expression of the F-actin probes affected invasion, we used DIC microscopy to assess basement membrane breaching beneath the anchor cell at the primary vulval precursor cell (1° VPC) four-cell stage, when invasion is expected to have occurred. All three probe strains exhibited 100% invasion, comparable to wild-type animals (**Figure 7B,C**).

**Figure 7:**
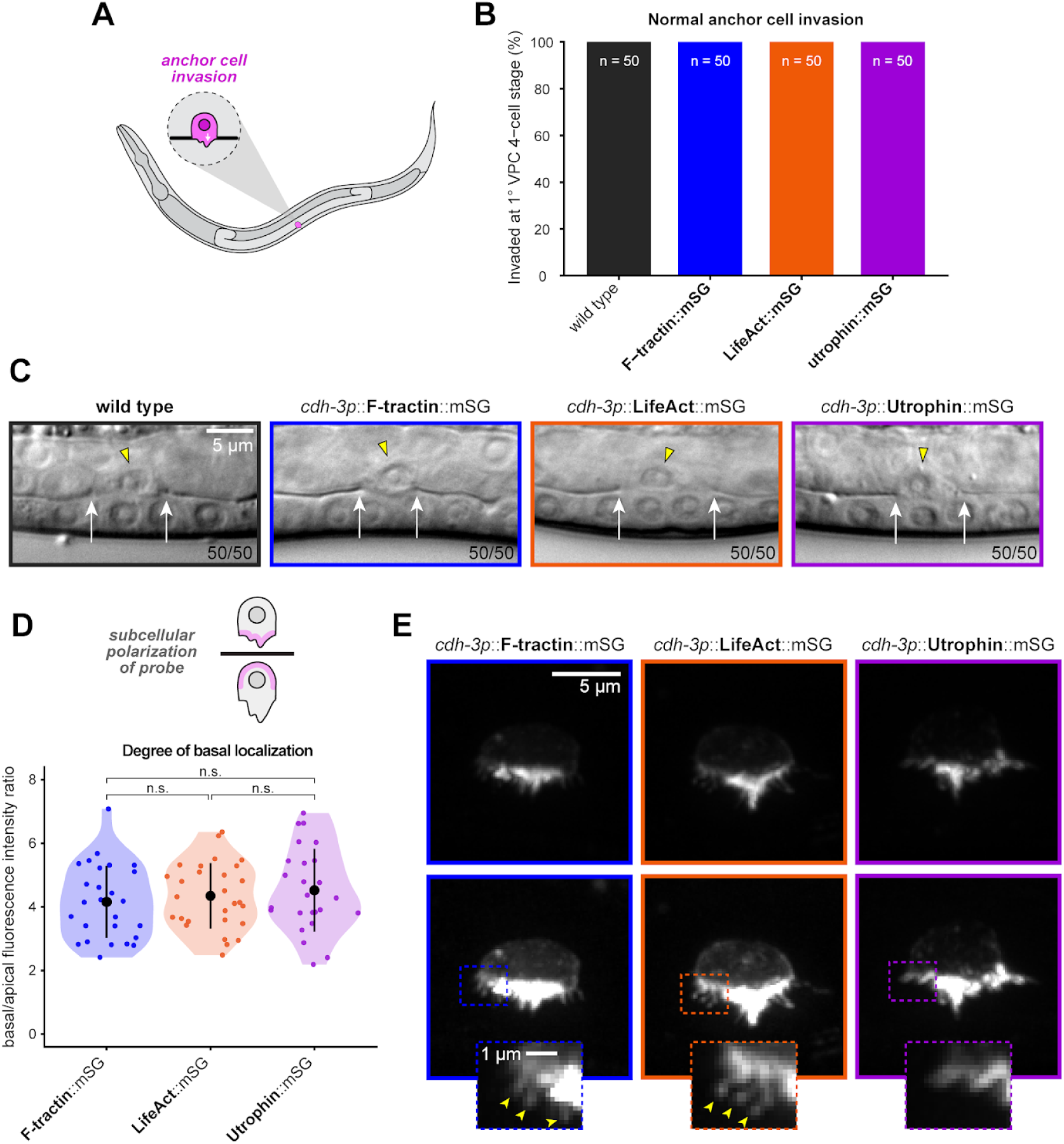
F-actin probes during anchor cell invasion. (A) Schematic depicting anchor cell invasion through the basement membrane. (B) Graph showing the percentage of animals expressing the indicated F-actin probes that exhibit normal anchor cell invasion, defined by the presence of a basement membrane gap by the four-cell stage of the primary vulval precursor cell (1° VPC). (C) Representative differential interference contrast (DIC) images of anchor cell invasion in the indicated strains. Yellow arrowheads indicate the anchor cell, and white arrows mark the boundaries of the basement membrane gap, visible as a loss of the phase-dense line. (D) Quantification of fluorescence asymmetry in the anchor cell, expressed as the ratio of basal to apical fluorescence intensity. Statistical significance was determined using Welch’s one-way ANOVA followed by Games-Howell post-hoc tests. (E) Representative micrographs of the anchor cell expressing the indicated F-actin probes under the *cdh-3* promoter. The bottom row is intentionally overexposed to highlight invadopodia, shown at higher magnification in the regions indicated by dashed boxes.

We next compared probe localization by measuring fluorescence intensities at the apical and basal cortices to determine the basal-to-apical ratio. The three strains exhibited no significant differences from one another, displaying a degree of basal F-actin enrichment consistent with previous findings (Ziel *et al*., 2009). All three probes revealed robust basal cortical fluorescence concentrated at the invadopodia (**Figure 7D,E**). However, enhancing image brightness to resolve weaker signals unveiled divergent labeling patterns: LifeAct and F-tractin clearly demarcated thin invasive protrusions, whereas fewer such structures were observed in Utrophin-expressing animals (**Figure 7E**). It remains unclear whether this difference reflects probe-specific labeling of F-actin or differences in the underlying actin structures caused by the probes. This discrepancy likely reflects a probe-specific labeling bias, as Utrophin CH-domain-containing markers preferentially bind stable, unbranched cortical actin networks in cultured cells and can underreport certain highly dynamic, transient protrusions captured by smaller peptides like LifeAct and F-tractin (Belin *et al*., 2014).

We next examined F-actin probe labeling in the sex myoblasts, which migrate during larval development to their final positions near the primary-fated vulval precursor cell P6.p before dividing to generate muscles required for egg laying (**Figure 8A**). To determine whether expression of the probes affected SM migration, we imaged the cells after they had completed migration but before their first division (**Figure 8B**). We assessed SM positioning by measuring the distance along the anterior–posterior body axis between the centers of the SM and P6.p nuclei. In all three probe strains, the sex myoblasts were positioned within approximately one cell length of P6.p, consistent with successful migration (**Figure 8C**).

**Figure 8:**
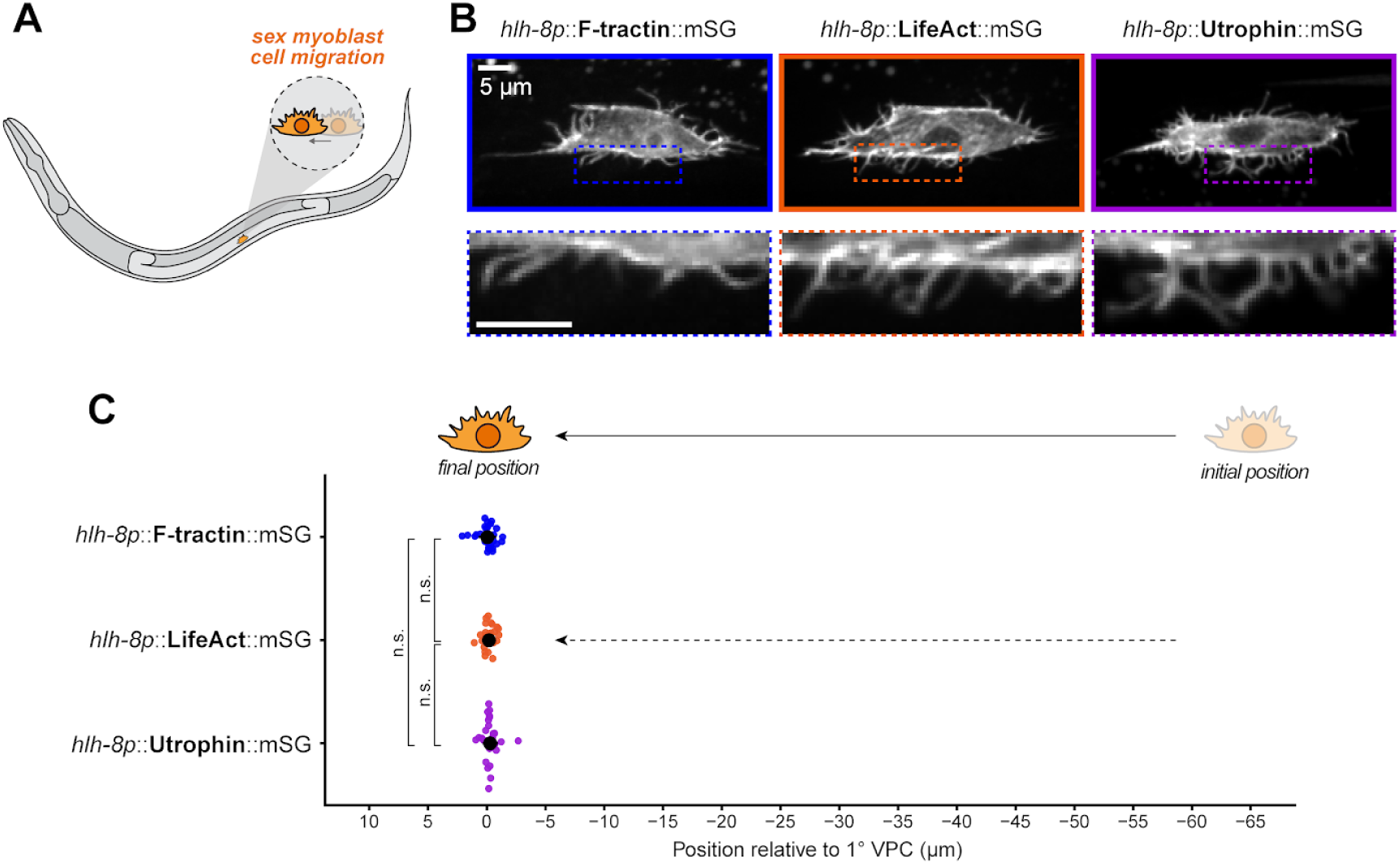
F-actin probes during sex myoblast migration. (A) Schematic depicting sex myoblast migration. (B) Representative micrographs of sex myoblasts expressing the indicated F-actin probes under the *hlh-8* promoter. Anterior is oriented to the left, and posterior to the right. Dashed boxes indicate regions shown at higher magnification to highlight filopodia. (C) Plot showing the final positions of sex myoblasts relative to the primary vulval precursor cell (1° VPC) in the indicated strains as a measure of migration success. Statistical significance was determined using Welch’s one-way ANOVA followed by Games-Howell post-hoc tests.

We also observed differences in the subcellular localization patterns of the probes in SMs. Filopodia extending from the sex myoblast were labeled by all three probes, with no consistent, obvious differences in their labeling patterns. F-tractin showed some nuclear enrichment, whereas Utrophin appeared excluded from the nucleus. In contrast, although LifeAct showed some nuclear localization at the four-cell stage, it was absent from the sex myoblast nucleus (**Figure 8B**). Although it remains unclear whether the apparent differences in cytoplasmic network density reflect probe-specific labeling or differences in the underlying actin organization of different cells, these observations highlight differences in the localization patterns of the probes across cellular contexts.

## Discussion

Actin filaments are fundamental to cell architecture and dynamics, and imaging of genetically encoded markers of F-actin makes it possible to see the distribution and dynamics of F-actin in living cells and organisms. Expression and/or imaging of some such markers has perturbed normal biological functions in some experimental systems, and has marked some F-actin-based structures less well than others in some systems. For this reason, systematic comparisons of F-actin markers in living cells or whole organisms can be valuable (Belin *et al*., 2013, 2014; Lemieux *et al*., 2014; Spracklen *et al*., 2014; Du *et al*., 2015). Here, we compared multiple markers *in vivo*, each inserted into the *C. elegans* genome as single copies. We examined a variety of cells that exhibit different F-actin based behaviors, including cytokinesis, apical constriction, production of filopodia, cell migration, and invasion through a basement membrane. Our new markers outperformed previously generated LifeAct-expressing strains in multiple respects. Comparisons among different F-actin markers revealed that LifeAct and Utrophin generally did as well as or outperformed F-tractin for brightly marking F-actin-containing structures while producing minimal or no disruption to the biological processes involved. Based on this work, we recommend certain strains for imaging specific processes in *C. elegans* (**Figure 9A**).

**Figure 9:**
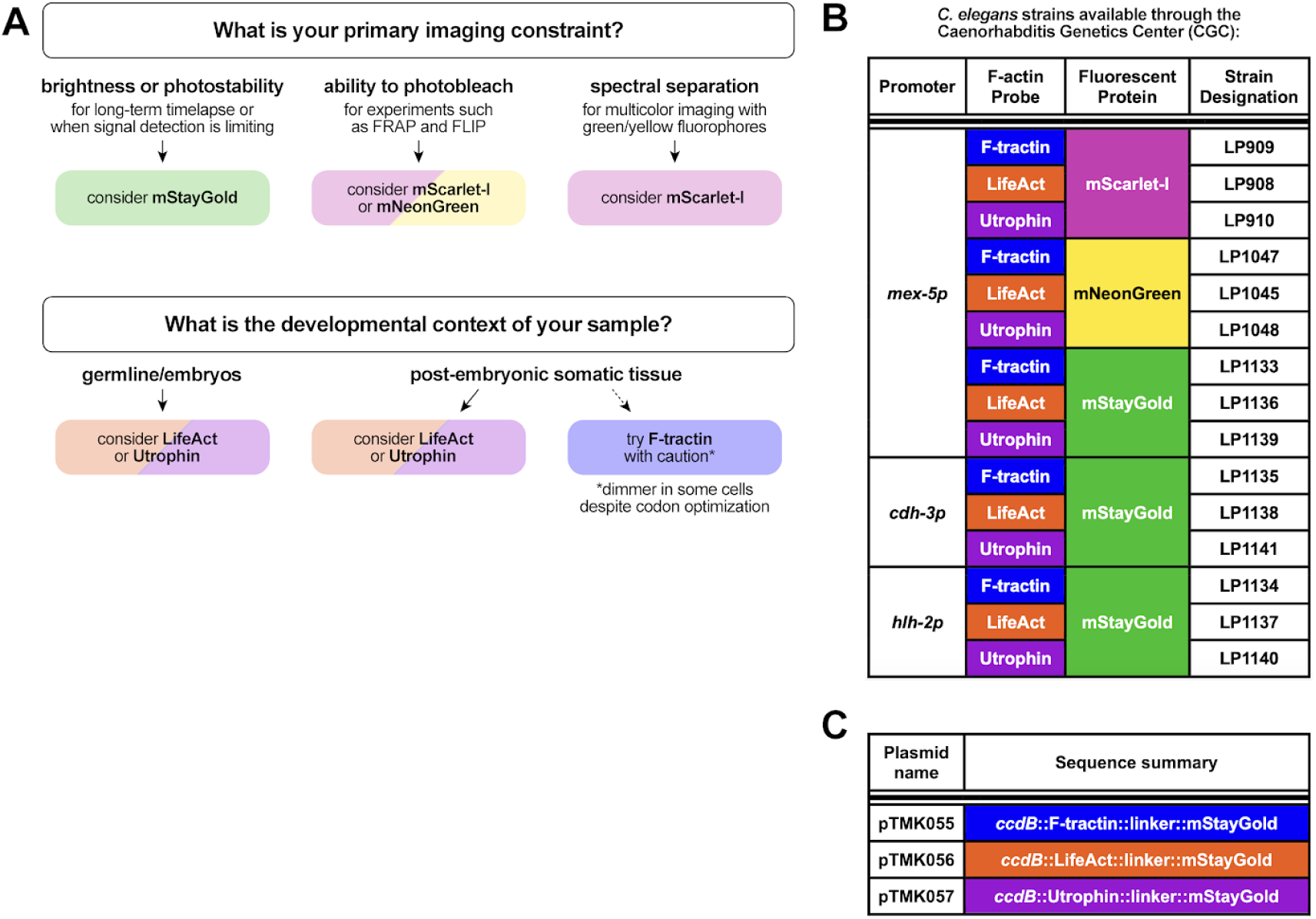
Resources for selecting and obtaining genetically encoded F-actin probes. (A) Guide to selecting F-actin probes based on imaging constraints, developmental context, and the structures to be visualized. (B) Table of strains generated in this study, available through the Caenorhabditis Genetics Center (CGC). (C) Plasmids available through Addgene. Each plasmid contains germline-optimized sequences encoding an F-actin probe and fluorescent protein separated by a linker. The fluorophore sequence is flanked by unique restriction sites, as is the promoter module, which contains a replaceable *ccdB* cassette that enables modular promoter insertion through negative selection.

We also compared different fluorescent proteins. Results were mostly as we had expected. mStayGold was a reliably bright and photostable fluorescent protein in the conditions we examined, consistent with the brightness and photostability of this monomeric variant of StayGold found previously (Ando *et al*., 2024). The photostability of mStayGold is a distinct advantage for imaging that requires extended illumination of samples, i.e. with many focal planes, at fine time resolution, and/or recording for a long time. Therefore mStayGold would be our first choice for fluorescently tagging a variety of proteins. mStayGold is not appropriate for experiments where photobleaching is desired, for example in fluorescence recovery after photobleaching (FRAP) experiments. mNeonGreen is generally a reliably bright fluorescent protein to use instead, although we have found previously that a subset of mNG-tagged proteins mislocalize to a pattern resembling some membranous organelles (Zhang *et al*., 2026 bioRxiv). We did not have that issue here with any of the F-actin markers tagged with mNeonGreen. Of course, FRAP of an actin-binding protein would need to be interpreted cautiously because FRAP results would depend on binding and unbinding of the marker to F-actin and not just actin turnover, unless markers can be demonstrated to bind quite stably to F-actin without perturbing its turnover. For imaging when green or yellow illumination of a separate marker is desired in combination with fluorescently marking F-actin, our results using mScarlet-I provide a validated alternative that can be used.

We found that the brightness of F-actin-containing structures varied to some degree among the three F-actin markers used, depending on the specific cells being marked. This variation might result from how well the markers access specific F-actin populations, consistent with previous experiments (Belin et al 2013, 2014), or it might result from differences in gene expression level, translation rates, or protein stability. These results suggest that rather than there being only a single F-actin marker that will work best in all cases, for some experiments testing more than one F-actin marker could be worthwhile, particularly when a first choice produces weak signal. As we had designed constructs for expression here in multiple contexts, the markers did not perturb most of the biological processes that we examined, suggesting that multiple choices are likely to be available for most cells and processes of interest.

Our study has some specific limitations. We have tested these constructs only in *C. elegans*, and their performance in different tissues varied, suggesting that probes’ performance will vary in other organisms as well. Furthermore, we tested only a few cell type-specific expression methods; stronger expression resulting from different strategies could perturb some biological processes. And although we tested our probes’ effects on a variety of F-actin-dependent behaviors, other F-actin-dependent events could potentially be perturbed even by the strategies that worked well here.

Our study provides a toolkit for examining F-actin dynamics *in vivo* in *C. elegans* in diverse cells and contexts (**Figure 9B,C**). We provide a standardized collection of single-copy, transgenic strains expressing in the germline F-tractin, LifeAct, and Utrophin with mNeonGreen, mStayGold, and mScarlet-I tags, which will be distributed by the Caenorhabditis Genetics Center. We also provide mStayGold strains for all three markers driving expression in the anchor cell or sex myoblasts. Data here provides baseline expectations for imaging each of the strains. We also provide modularly designed plasmids, to be distributed via AddGene, that can be modified for expression in other cells or tissues, and for swapping in other fluorescent proteins, actin-markers, and/or proteins.

## Materials and Methods

### *C. elegans* maintenance

*C. elegans* were maintained on nematode growth medium (NGM) plates seeded with *Escherichia coli* OP50, as previously described (Brenner, 1974; Stiernagle, 2006). The strains used in this study are listed in **Table 1**.

**Table 1.**
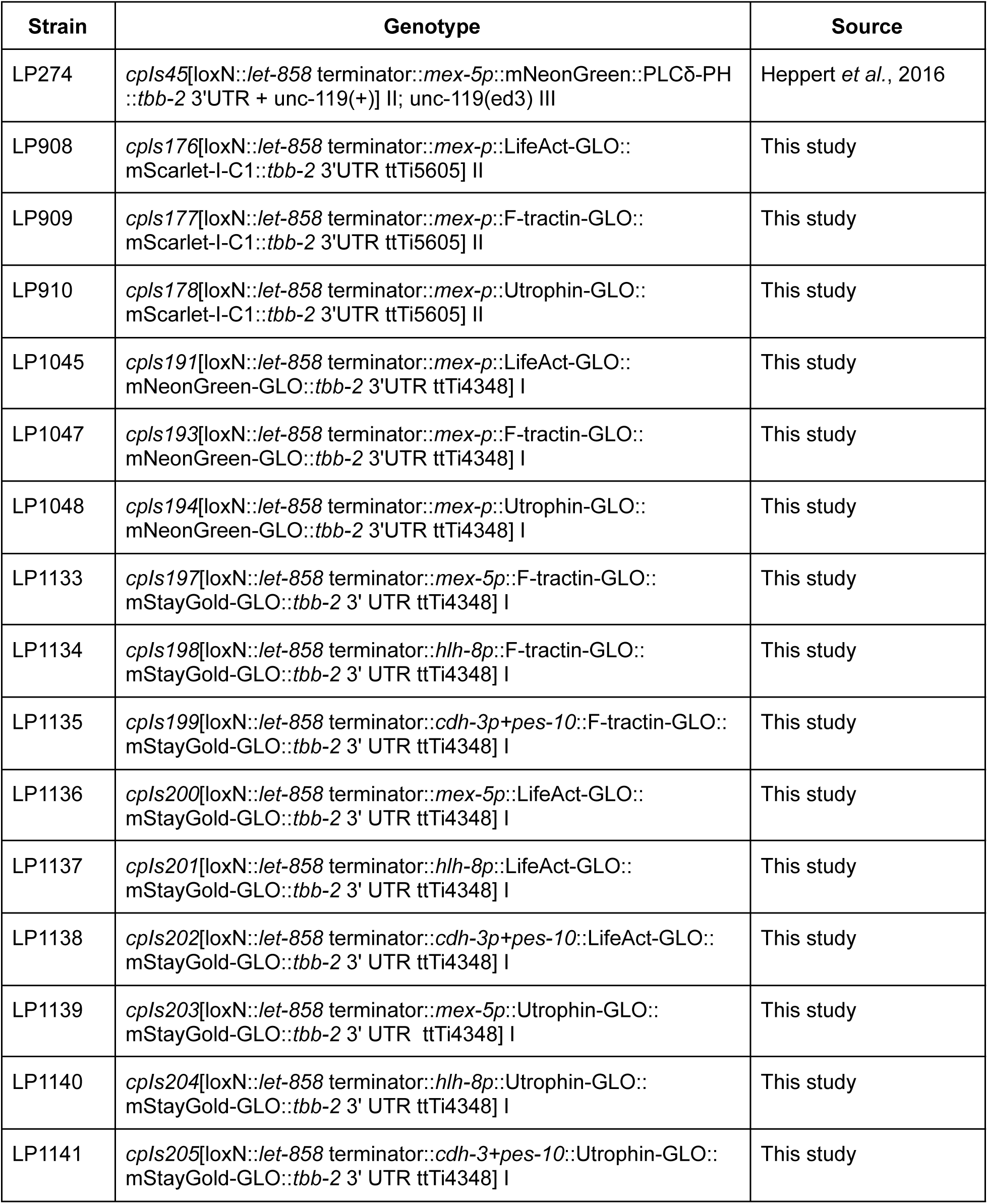
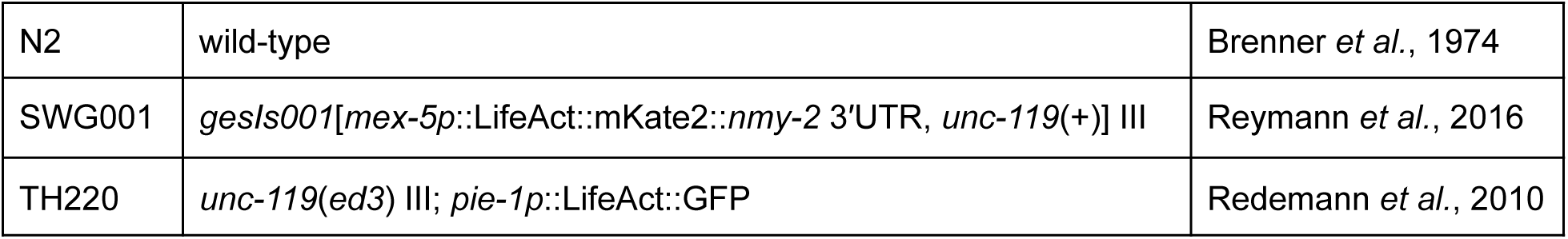
Strains used in this study.

| Strain | Genotype | Source |
|---|---|---|
| LP274 | *cpIs45*[loxN::*let-858* terminator::*mex-5p*::mNeonGreen::PLCδ-PH::*tbb-2* 3'UTR + unc-119(+)] II; unc-119(ed3) III | Heppert *et al.*, 2016 |
| LP908 | *cpIs176*[loxN::*let-858* terminator::*mex-p*::LifeAct-GLO::mScarlet-I-C1::*tbb-2* 3'UTR ttTi5605] II | This study |
| LP909 | *cpIs177*[loxN::*let-858* terminator::*mex-p*::F-tractin-GLO::mScarlet-I-C1::*tbb-2* 3'UTR ttTi5605] II | This study |
| LP910 | *cpIs178*[loxN::*let-858* terminator::*mex-p*::Utrophin-GLO::mScarlet-I-C1::*tbb-2* 3'UTR ttTi5605] II | This study |
| LP1045 | *cpIs191*[loxN::*let-858* terminator::*mex-p*::LifeAct-GLO::mNeonGreen-GLO::*tbb-2* 3'UTR ttTi4348] I | This study |
| LP1047 | *cpIs193*[loxN::*let-858* terminator::*mex-p*::F-tractin-GLO::mNeonGreen-GLO::*tbb-2* 3'UTR ttTi4348] I | This study |
| LP1048 | *cpIs194*[loxN::*let-858* terminator::*mex-p*::Utrophin-GLO::mNeonGreen-GLO::*tbb-2* 3'UTR ttTi4348] I | This study |
| LP1133 | *cpIs197*[loxN::*let-858* terminator::*mex-5p*::F-tractin-GLO::mStayGold-GLO::*tbb-2* 3' UTR ttTi4348] I | This study |
| LP1134 | *cpIs198*[loxN::*let-858* terminator::*hlh-8p*::F-tractin-GLO::mStayGold-GLO::*tbb-2* 3' UTR ttTi4348] I | This study |
| LP1135 | *cpIs199*[loxN::*let-858* terminator::*cdh-3p+pes-10*::F-tractin-GLO::mStayGold-GLO::*tbb-2* 3' UTR ttTi4348] I | This study |
| LP1136 | *cpIs200*[loxN::*let-858* terminator::*mex-5p*::LifeAct-GLO::mStayGold-GLO::*tbb-2* 3' UTR ttTi4348] I | This study |
| LP1137 | *cpIs201*[loxN::*let-858* terminator::*hlh-8p*::LifeAct-GLO::mStayGold-GLO::*tbb-2* 3' UTR ttTi4348] I | This study |
| LP1138 | *cpIs202*[loxN::*let-858* terminator::*cdh-3p+pes-10*::LifeAct-GLO::mStayGold-GLO::*tbb-2* 3' UTR ttTi4348] I | This study |
| LP1139 | *cpIs203*[loxN::*let-858* terminator::*mex-5p*::Utrophin-GLO::mStayGold-GLO::*tbb-2* 3' UTR ttTi4348] I | This study |
| LP1140 | *cpIs204*[loxN::*let-858* terminator::*hlh-8p*::Utrophin-GLO::mStayGold-GLO::*tbb-2* 3' UTR ttTi4348] I | This study |
| LP1141 | *cpIs205*[loxN::*let-858* terminator::*cdh-3+pes-10*::Utrophin-GLO::mStayGold-GLO::*tbb-2* 3' UTR ttTi4348] I | This study |
| N2 | wild-type | Brenner *et al.*, 1974 |
| SWG001 | *gesIs001*[*mex-5p*::LifeAct::mKate2::*nmy-2* 3′UTR, *unc-119*(+)] III | Reymann *et al.*, 2016 |
| TH220 | *unc-119*(*ed3*) III; *pie-1p*::LifeAct::GFP | Redemann *et al.*, 2010 |

### Strain generation

The genetically encoded F-actin probes were integrated as single-copy transgenes at defined genomic integration sites: ttTi4348 on chromosome I for mNeonGreen and mStayGold constructs, and ttTi5605 on chromosome II for mScarlet-I constructs. Repair templates were generated using plasmids containing the 5′ and 3′ homology arms corresponding to each integration site, along with a self-excising cassette (SEC) that enabled initial selection and subsequent homozygosing based on hygromycin resistance and the roller phenotype (Dickinson *et al*., 2015).

Previously generated codon-optimized (C1) mScarlet-I and germline-optimized mNeonGreen sequences were amplified from existing plasmids and used in construct assembly. The mStayGold and F-actin probe sequences (F-tractin, LifeAct, and Utrophin) were adapted using a previously published germline-optimization algorithm (Fielmich *et al*., 2018) and modified to remove piRNA homology (Priyadarshini et al., 2022). These sequences were ordered as gene fragments from IDT or Twist Bioscience.

Plasmids were assembled using restriction digestion of the repair template backbone, amplification of inserts, and Gibson assembly. For the mStayGold constructs, a ccdB site was first inserted as a selectable placeholder for subsequent promoter insertion. Unique restriction sites (NotI, AgeI, NgoMIV, and AflII) were also incorporated to facilitate modular exchange of promoter and fluorescent protein sequences. Repair templates were co-injected with plasmids encoding ubiquitously expressed Cas9 and a single-guide RNA (sgRNA) targeting the intended genomic integration site. Resulting strains were made homozygous, floxed (to remove selection cassette), and sequence-verified. The plasmids and DNA sequences used in this study are listed in **Tables 2 and 3**, respectively.

**Table 2.** Plasmids.

| Plasmid | Description | Source |
|---|---|---|
| pAP082 | *eft-3p*::Cas9 + ttTi4348 (chromosome I) sgRNA | Pani and Goldstein, 2018 |
| pDD122 | *eft-3p*::Cas9 + ttTi5605 (chromosome II) sgRNA | Dickinson *et al.*, 2013 |
| pTMK055 | loxN::SEC::loxN::*let-858* terminator::*ccdB*::<br>F-tractin-GLO::linker::mStayGold-GLO::*tbb-2* 3' UTR<br>repair template for ttTi4348 locus | This study |
| pTMK056 | loxN::SEC::loxN::*let-858* terminator::*ccdB*::<br>LifeAct-GLO::linker::mStayGold-GLO::*tbb-2* 3' UTR<br>repair template for ttTi4348 locus | This study |
| pTMK057 | loxN::SEC::loxN::*let-858* terminator::*ccdB*::<br>Utrophin-GLO::linker::mStayGold-GLO::*tbb-2* 3' UTR<br>repair template for ttTi4348 locus | This study |
| pTMK058 | loxN::SEC::loxN::*let-858* terminator::*mex-5p*::<br>F-tractin-GLO::linker::mStayGold-GLO::*tbb-2* 3' UTR<br>repair template for ttTi4348 locus | This study |
| pTMK059 | loxN::SEC::loxN::*let-858* terminator::*hlh-8p*::<br>F-tractin-GLO::linker::mStayGold-GLO::*tbb-2* 3' UTR<br>repair template for ttTi4348 locus | This study |
| pTMK060 | loxN::SEC::loxN::*let-858* terminator::*cdh-3p+pes-10p*::<br>F-tractin::linker::mStayGold-GLO::*tbb-2* 3' UTR<br>repair template for ttTi4348 locus | This study |
| pTMK061 | loxN::SEC::loxN::*let-858* terminator::*mex-5p*::<br>LifeAct-GLO::linker::mStayGold-GLO::*tbb-2* 3' UTR<br>repair template for ttTi4348 locus | This study |
| pTMK062 | loxN::SEC::loxN::*let-858* terminator::*hlh-8p*::<br>LifeAct-GLO::linker::mStayGold-GLO::*tbb-2* 3' UTR<br>repair template for ttTi4348 locus | This study |
| pTMK063 | loxN::SEC::loxN::*let-858* terminator::*cdh-3p+pes-10p*::<br>LifeAct::linker::mStayGold-GLO::*tbb-2* 3' UTR<br>repair template for ttTi4348 locus | This study |
| pTMK064 | loxN::SEC::loxN::*let-858* terminator::*mex-5p*::<br>Utrophin-GLO::linker::mStayGold-GLO::*tbb-2* 3' UTR<br>repair template for ttTi4348 locus | This study |
| pTMK065 | loxN::SEC::loxN::*let-858* terminator::*hlh-8p*::<br>Utrophin-GLO::linker::mStayGold-GLO::*tbb-2* 3' UTR<br>repair template for ttTi4348 locus | This study |
| pTMK066 | loxN::SEC::loxN::*let-858* terminator::*cdh-3p+pes-10p*::<br>Utrophin::linker::mStayGold-GLO::*tbb-2* 3' UTR<br>repair template for ttTi4348 locus | This study |
| pPZ013 | loxN::SEC::loxN::*let-858* terminator::*mex-5p*::<br>LifeAct-GLO::linker::mScarlet-I-C1::*tbb-2* 3' UTR<br>repair template for ttTi5605 locus | This study |
| pPZ014 | loxN::SEC::loxN::*let-858* terminator::*mex-5p*::<br>F-tractin-GLO::linker::mScarlet-I-C1::*tbb-2* 3' UTR<br>repair template for ttTi5605 locus | This study |
| pPZ015 | loxN::SEC::loxN::*let-858* terminator::*mex-5p*::<br>Utrophin-GLO::linker::mScarlet-I-C1::*tbb-2* 3' UTR<br>repair template for ttTi4348 locus | This study |
| pPZ016 | loxN::SEC::loxN::*let-858* terminator::*mex-5p*::<br>LifeAct-GLO::linker::mNeonGreen-GLO::*tbb-2* 3' UTR<br>repair template for ttTi4348 locus | This study |
| pPZ023 | loxN::SEC::loxN::*let-858* terminator::*mex-5p*::<br>F-tractin-GLO::linker::mNeonGreen-GLO::*tbb-2* 3' UTR<br>repair template for ttTi4348 locus | This study |
| pPZ024 | loxN::SEC::loxN::*let-858* terminator::*mex-5p*::<br>Utrophin-GLO::linker::mNeonGreen-GLO::*tbb-2* 3' UTR<br>repair template for ttTi4348 locus | This study |

**Table 3.**
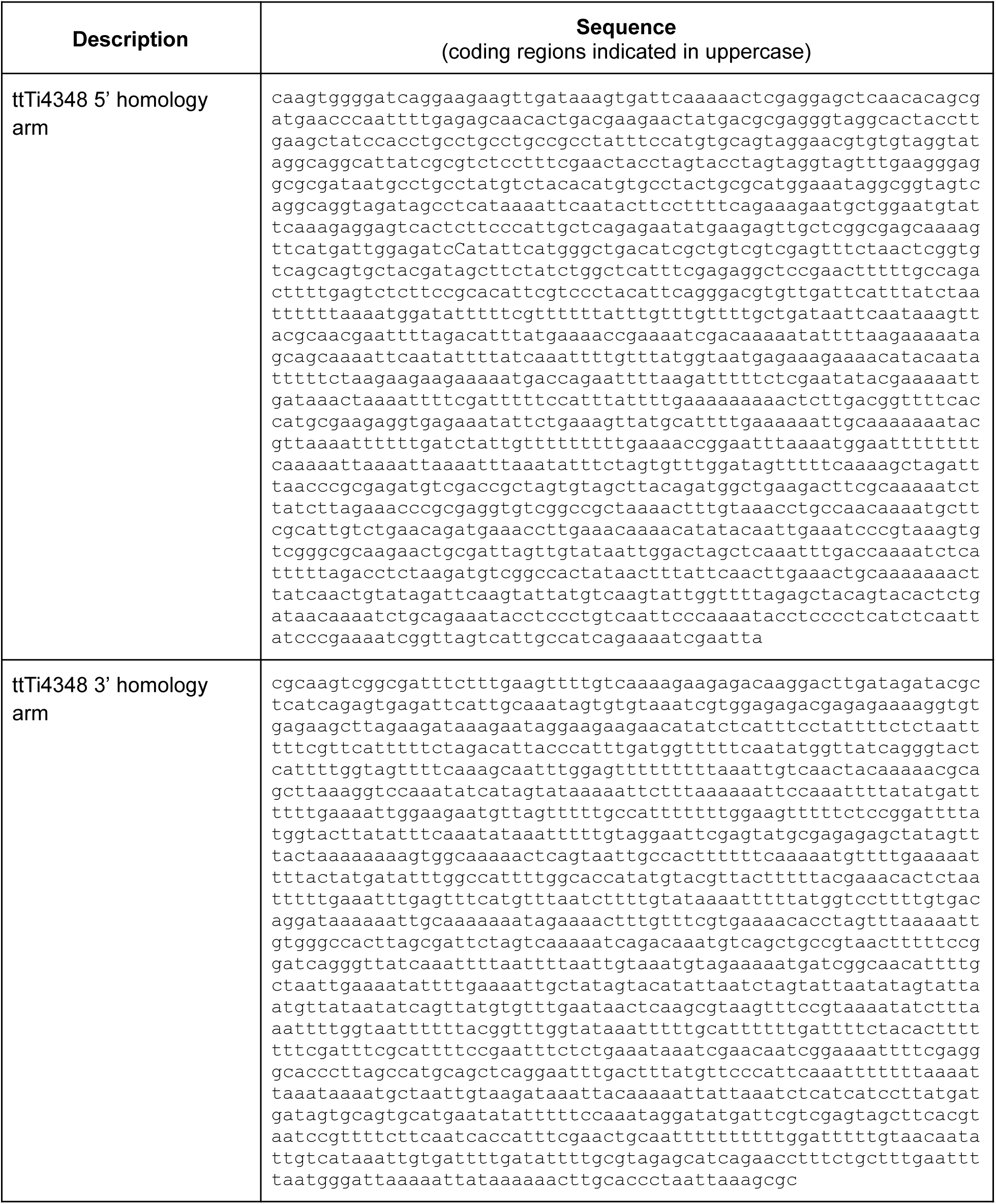
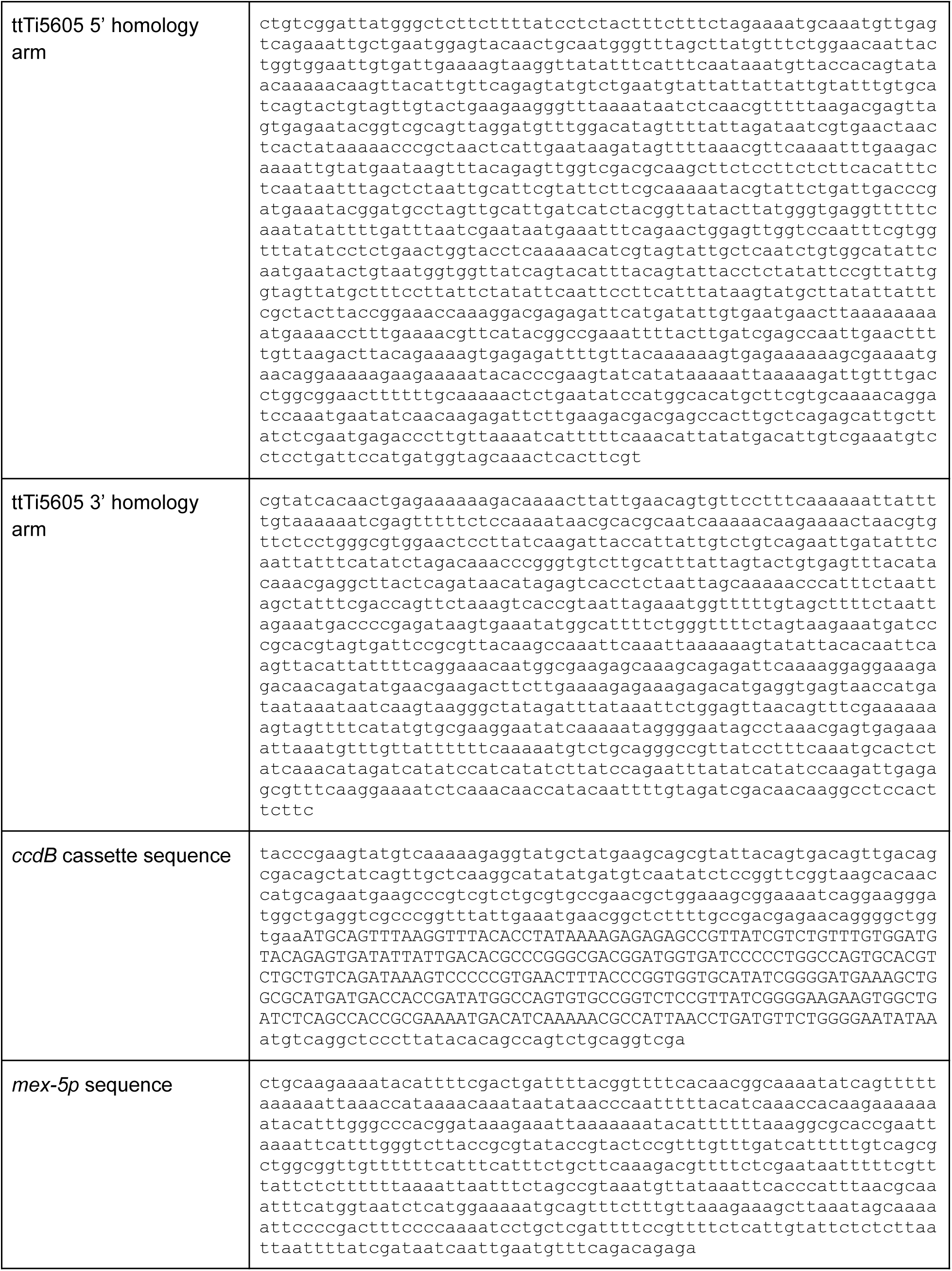
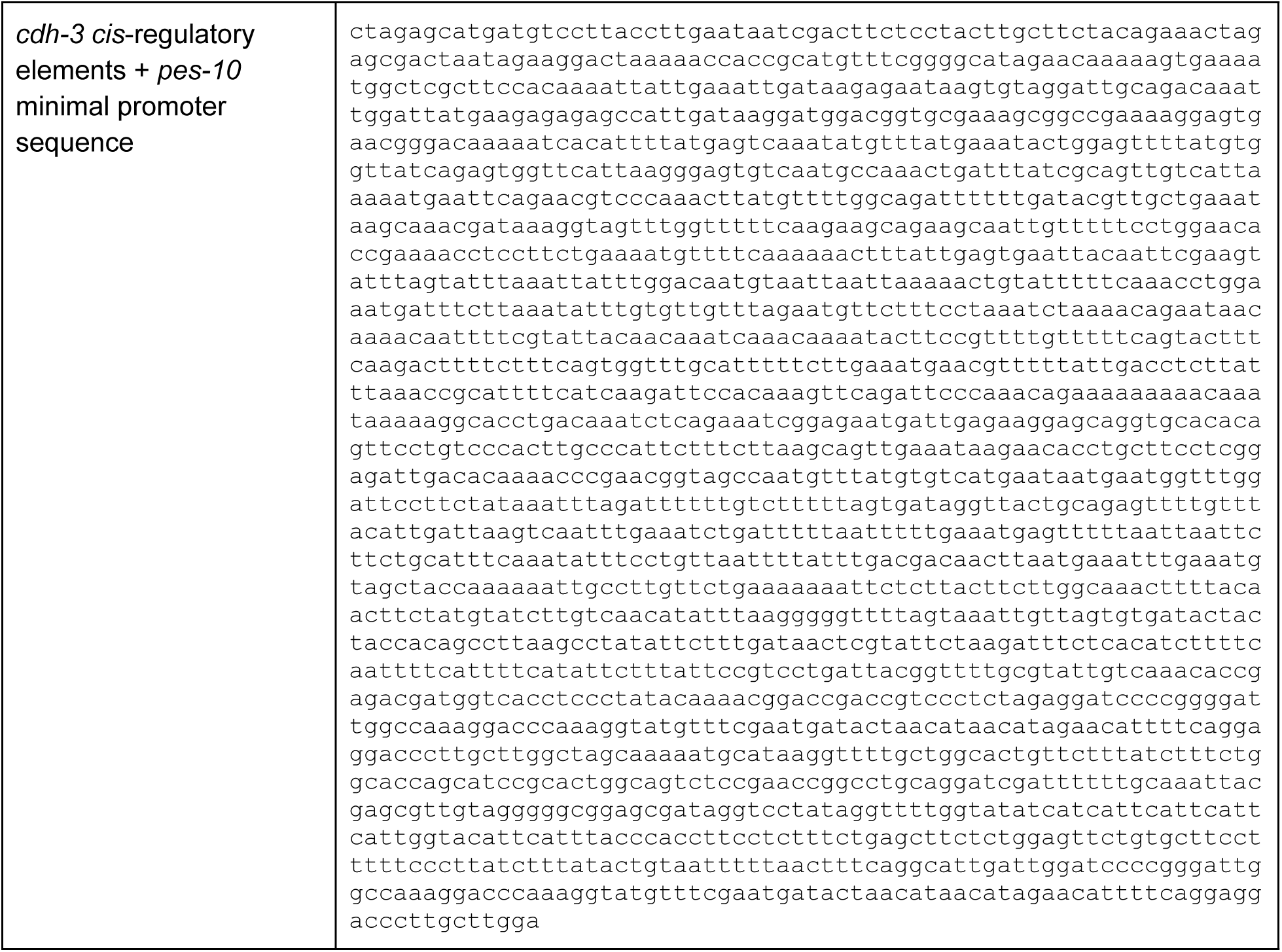
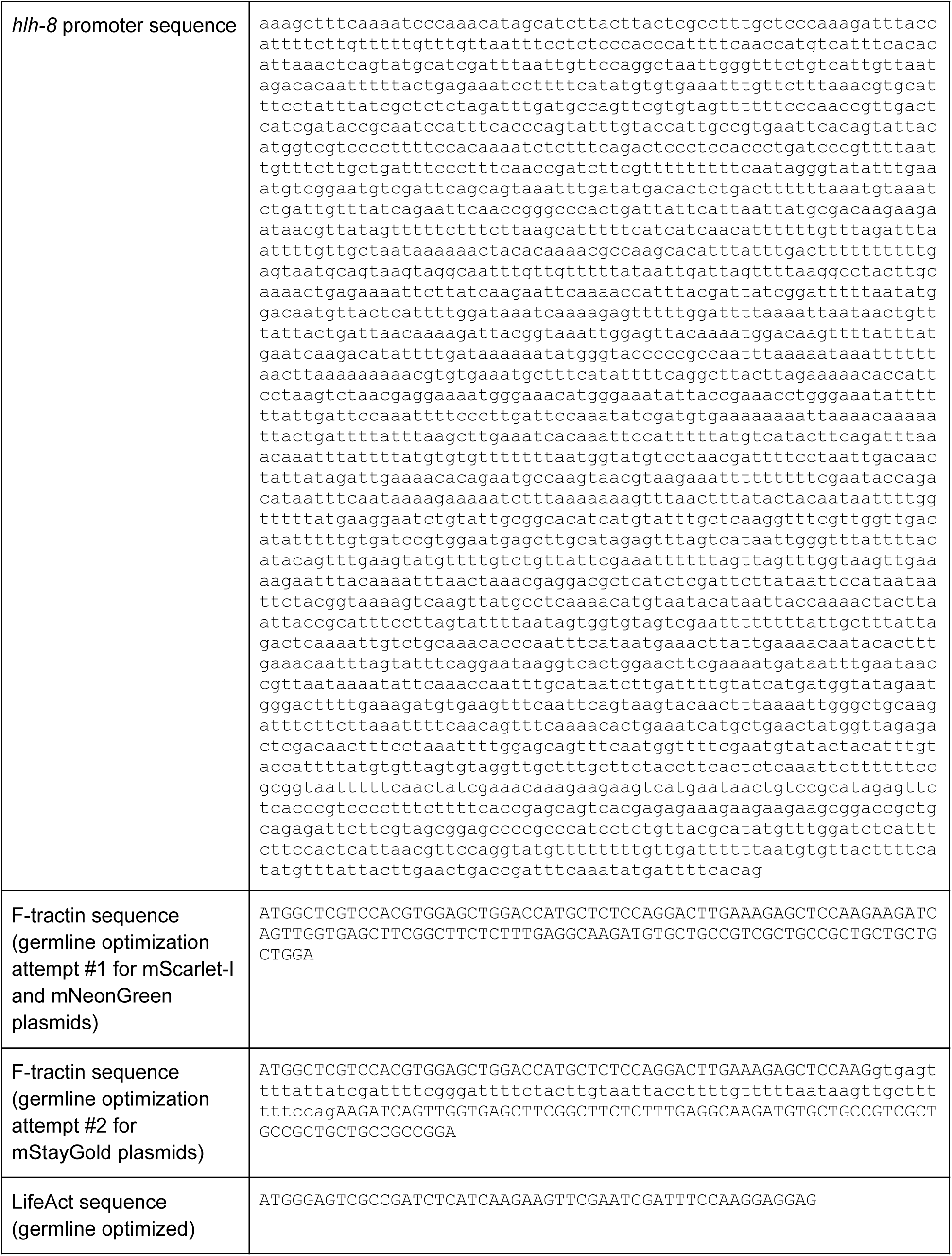
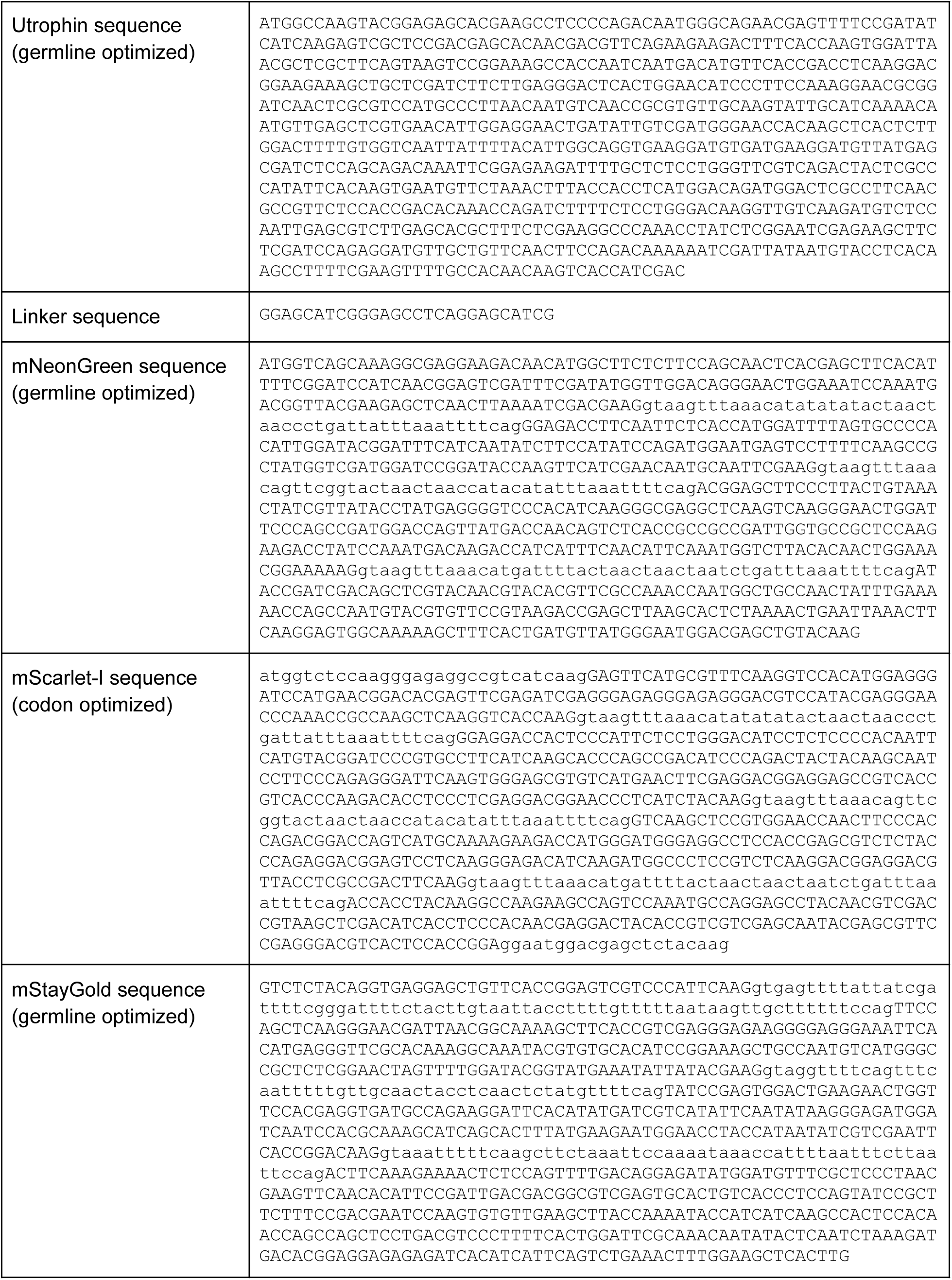
Sequences used in this study.

### Brood size and embryonic lethality measurements

Brood size and embryonic lethality assays were conducted in accordance with a previous protocol (Kwah and Jaramillo-Lambert, 2023). Briefly, L4 stage larvae were singled out onto NGM plates. Worms were transferred to new plates every 24 hours for four times to capture all offspring produced within a 96 hour period. All plates were kept in a 20°C incubator. All worms and unhatched fertilized embryos on a plate were counted 48 hour post worm transfer onto the plate. Any unhatched fertilized embryos were assumed to be unviable. Brood size was calculated as the total number of worms and fertilized embryos. Embryonic lethality was calculated as the proportion of the total brood that did not hatch during the observation period.

### Body length measurement

Worms were synchronized by picking 10 gravid hermaphrodites onto an NGM plate, allowed to lay eggs, and removed after 2 hours. Worm body lengths were measured 48, 72, and 96 hours post egg lay. Image acquisition was performed using the Adonstar AD246S digital microscope with a photo resolution of 4032×3024 pixels. Lens A was used, and it has a working distance of 12-320mm, at 720× magnification with 3x digital zoom. Images were uploaded onto a custom Desmos graph (https://www.desmos.com/calculator/yhbu9reod2) that calculates the length of a degree 8 Bézier curve that can be manipulated with 9 control points. The length calculation was calibrated by an image of a microscope micrometer ruler. After calibration, worm body lengths were acquired upon fitting the Bézier curve to worm images.

### Fluorescence and differential interference contrast microscopy

With the exception of cytokinesis experiments, fluorescence imaging was performed using a Hamamatsu C11440-22CU ORCA Flash 4.0 sCMOS camera mounted on a Nikon Eclipse Ti inverted microscope equipped with a Yokogawa QLC100/CSU10 spinning-disk scan head. Images were acquired using a 60× Plan Apo λ oil-immersion objective (NA 1.4) with a 1.5× magnifier, except for sex myoblast imaging, which was performed using a 40× Plan Fluor oil-immersion objective (NA 1.3). Excitation was provided by 488-nm and 561-nm lasers.

DIC imaging for developmental staging, assessment of anchor cell invasion, and timing of gastrulation was performed using a Nikon Eclipse 800 microscope equipped with a Diagnostic Instruments SPOT2 camera and a 60× oil-immersion objective (NA 1.4).

### Preparing live samples

For imaging four-cell-stage embryos and embryos undergoing apical constriction, embryos were dissected from gravid hermaphrodites and mounted on slides in *C. elegans* egg buffer. Coverslips were sealed with VALAP. For ventral mounts, approximately 23-µm glass beads (Whitehouse Scientific Monodisperse Standards, MS0023) and clay feet were used to separate the slide and coverslip. For lateral mounts, embryos were mounted on 2.5% agar pads.

For imaging post-embryonic development, animals were mounted on 2.5% agar pads in a drop of M9 buffer containing 20 mM sodium azide for immobilization.

### Fluorescence intensity and photobleaching measurements

Fluorescence intensity was quantified using FIJI. Strains expressing green fluorescent proteins were imaged under identical acquisition settings, as were strains expressing red fluorescent proteins. For four-cell-stage embryos, fluorescence intensity was measured by selecting the image plane corresponding to the embryo midplane, drawing a region of interest (ROI) around the entire embryo, and measuring the mean gray value of the enclosed pixels. The same approach was used to measure fluorescence intensity in anchor cells and sex myoblasts, except that ROIs were drawn around the individual cells. With the exception of four-cell-stage embryo measurements, for which wild-type embryos were imaged to establish a reference for autofluorescence, background fluorescence was subtracted by measuring the mean gray value of an ROI outside the sample to account for camera background. Photobleaching was assessed by continuously illuminating samples at 100% laser power for 120 seconds using 488-nm excitation for GFP, mNeonGreen, and mStayGold, or 561-nm excitation for mKate2 and mScarlet-I. Mean gray values were measured as described above and normalized to the initial fluorescence intensity. Normalized fluorescence intensity was plotted over time, and fluorescence decay constants were determined by fitting the data to an exponential decay curve.

### Developmental staging

Embryos were collected from gravid adults by hypochlorite treatment (Porta-de-la-Riva *et al*., 2012) and allowed to hatch in M9 solution without food for 24 hours to obtain synchronized L1-arrested larvae. Larvae were then plated and allowed to develop for 39 hours at 20°C. DIC imaging was used to classify developmental substages. L3 substages were classified according to the number of primary vulval precursor cells (1° VPCs), whereas L4 substages were determined based on vulval morphology using an established staging system (Mok *et al*., 2015).

### Cytokinetic ring closure time measurements

Sample preparation and live imaging of embryos were performed at 20°C. Gravid hermaphrodites were dissected in a droplet of M9 on the coverglass and mounted onto a 2% agarose pad. Acquisition was performed on a Nikon A1R microscope body with a Gallium arsenide phosphide photo-multiplier tube detector using a 60×1.27 NA Nikon Water Immersion Objective. The cytokinetic ring closure time was obtained by determining the time elapsed between initial furrow ingression in the mid-plane and the meeting of the two tips of the ingressing cleavage furrow in the z-plane with the widest gap. Image processing and analysis were performed using Fiji (ImageJ; National Institutes of Health).

### Polarity measurements

Fluorescence polarization between the AB and P_1_ cells was quantified by drawing separate regions of interest (ROIs) around each cell and measuring the mean gray value within each ROI. The AB-to-P_1_ fluorescence ratio was calculated by dividing the mean gray value of AB by that of P1.

For anchor cell polarity, a 2-pixel-wide ROI was drawn along the basal and apical cortices, and the mean gray value was measured for each region. The basal-to-apical fluorescence ratio was calculated by dividing the mean gray value of the basal cortex by that of the apical cortex.

### Phalloidin staining

Embryos were isolated from gravid hermaphrodites by hypochlorite treatment and fixed for 15 min at room temperature in a solution containing 60 mM PIPES, 10 mM EGTA, 25 mM HEPES, 1 mM MgCl₂, 4% formaldehyde, 0.2% glutaraldehyde, and 0.1 mg/mL lysolecithin. Fixed embryos were permeabilized and washed in PBS containing 0.1% Triton X-100, then incubated with Alexa Fluor 488 phalloidin in PBS containing 0.1% Triton X-100 for 1 hour at room temperature. Embryos were subsequently washed three times in PBS and mounted for imaging.

### Imaging apical constriction

Differential interference contrast images were acquired at 1-minute intervals across multiple focal planes. The duration of apical constriction was measured from MSx division until both Ea and Ep were fully internalized beneath the embryo surface and completely covered by neighboring cells. For fluorescence imaging, embryos were imaged every 2-3 minutes.

### Imaging anchor cells and sex myoblasts

Animals were synchronized by hypochlorite treatment and L1 arrest as described above. Following plating, animals were allowed to develop for 24 hours at 25°C for anchor cell imaging or 26 hours at 20°C for sex myoblast imaging. Anchor cell invasion was assessed by DIC microscopy, with loss of the phase-dense line beneath the anchor cell, representing the basement membrane, indicating successful invasion. Anchor cells and sex myoblasts were imaged by fluorescence microscopy using identical acquisition settings within each cell type.

### Data visualization and statistical analysis

Molecular structure visualizations in Figure 1A were generated using PyMOL Molecular Graphics System (Schrödinger, LLC). Representative micrographs were processed in ImageJ/FIJI (Schindelin *et al*., 2012). Statistical analyses and data visualization were performed using R (Wickham, 2016). Statistical tests used for individual experiments are indicated in the corresponding figure legends. Statistical significance is denoted as follows: n.s., not significant; * *p* ≤ 0.05; ** *p* ≤ 0.01; *** *p* ≤ 0.001. Adobe Illustrator was used to create cell cartoons and assemble figures.

## Acknowledgments

We thank Dave Sherwood for sharing the sequence containing the *cdh-3* cis-regulatory sequence used to drive anchor cell-specific expression. We also thank the Munro laboratory for sharing their phalloidin staining protocol. This work was supported by the National Institutes of Health (R35GM134838 to B.G. and R35GM144238 to A.S.B.), National Science Foundation (MCB 2153790 to A.S.B.), American Cancer Society (A24-0591-001 to T.N.M.-K.), and L’Oréal For Women in Science Program (to T.N.M.-K.).

## Abbreviations

DIC: differential interference contrast
mK2: mKate2
mNG: mNeonGreen
mSc: mScarlet-I
mSG: mStayGold

## Figure legends

**Supplemental Figure S1.**
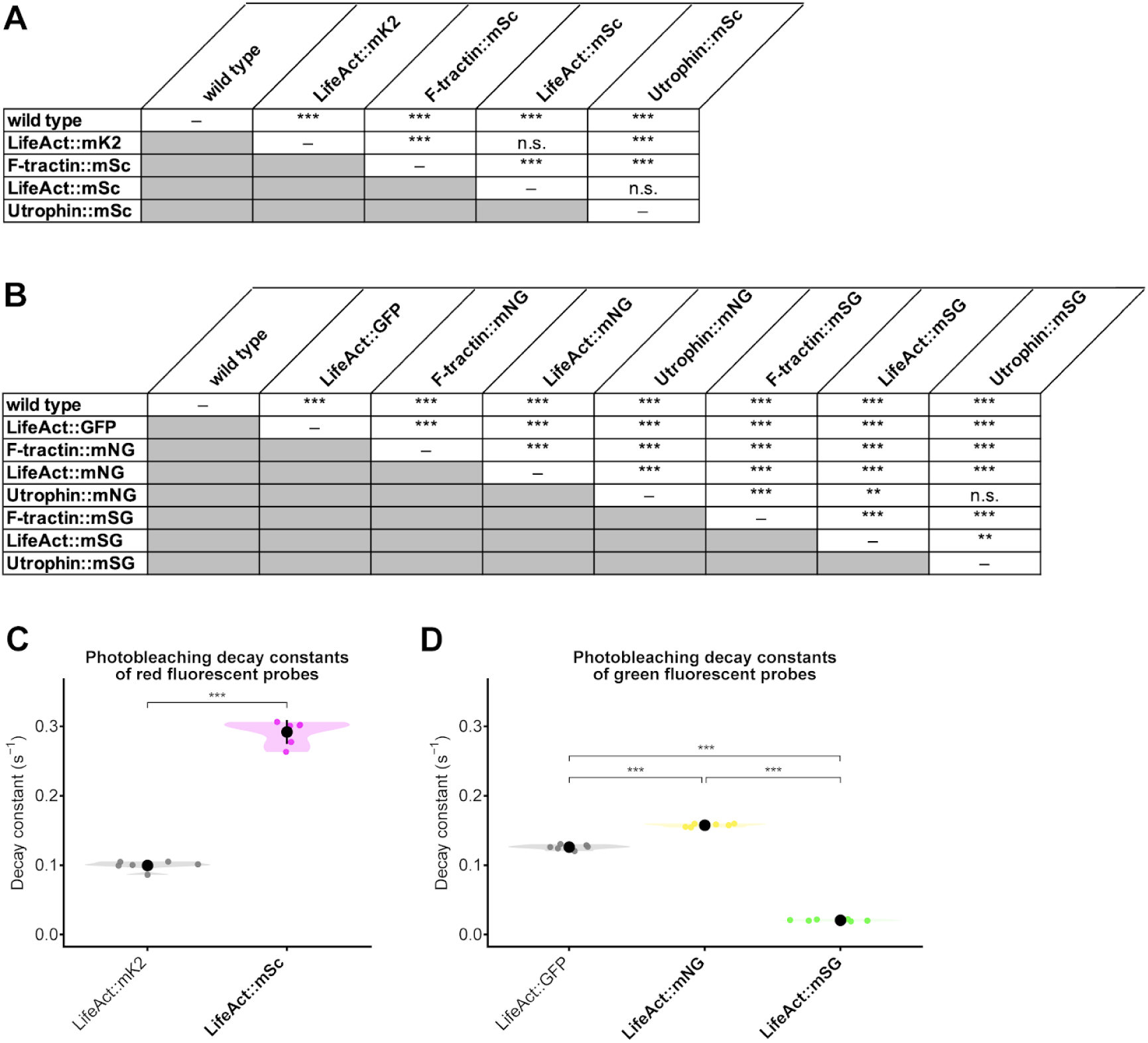
Statistical comparisons of fluorescence intensity and photostability among F-actin probe strains. (A) Pairwise comparisons among wild type animals, the previously established *mex-5p*::LifeAct::mKate2 strain, and three newly generated mScarlet-I-tagged F-actin probe strains. (B) Pairwise comparisons among wild type animals, the previously established*pie-1p*::LifeAct::GFP strain, and six newly generated mNeonGreen- or mStayGold-tagged F-actin probe strains. (C-D) Comparison of photobleaching decay constants (*k*) among red (C) and green (D) fluorescent F-actin probes. Higher decay constants indicate faster photobleaching and lower photostability. Statistical significance was assessed using Welch’s one-way ANOVA followed by Games-Howell post-hoc tests (A, B), Welch’s unpaired *t*-test (C), or pairwise Welch’s *t*-tests with Holm correction for multiple comparisons (D).

**Supplemental Figure S2.**
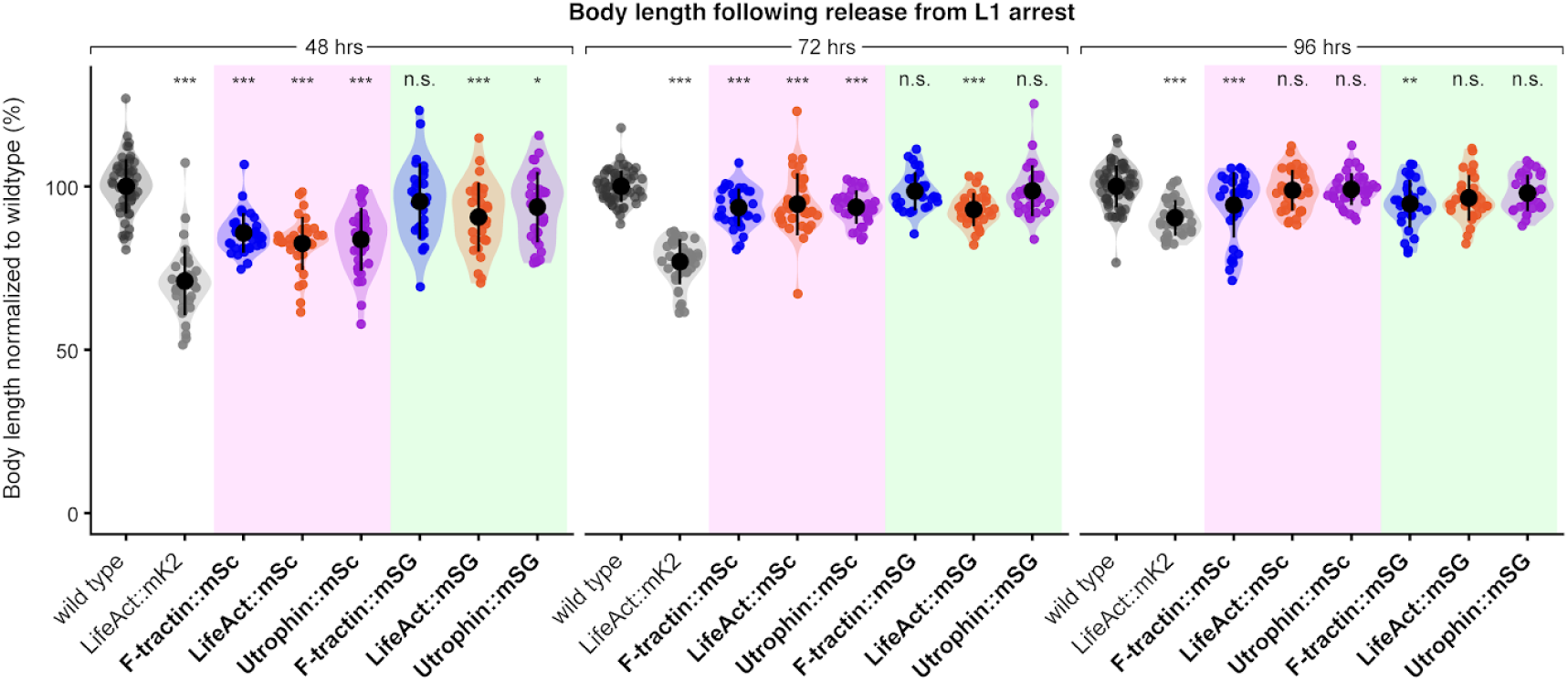
Body length of animals expressing fluorescently tagged F-actin probes. Body length of animals expressing newly generated mScarlet-I- and mStayGold-tagged F-actin probes compared with the established *mex-5p*::LifeAct::mKate2 probe at 48, 72, and 96 hours after release from L1 arrest. Body length is normalized to the mean body length of wild-type animals at each time point. Statistical significance was determined separately at each time point using one-way ANOVA followed by Dunnett’s post-hoc test, comparing each strain to wild type.

**Supplemental Figure S3.**
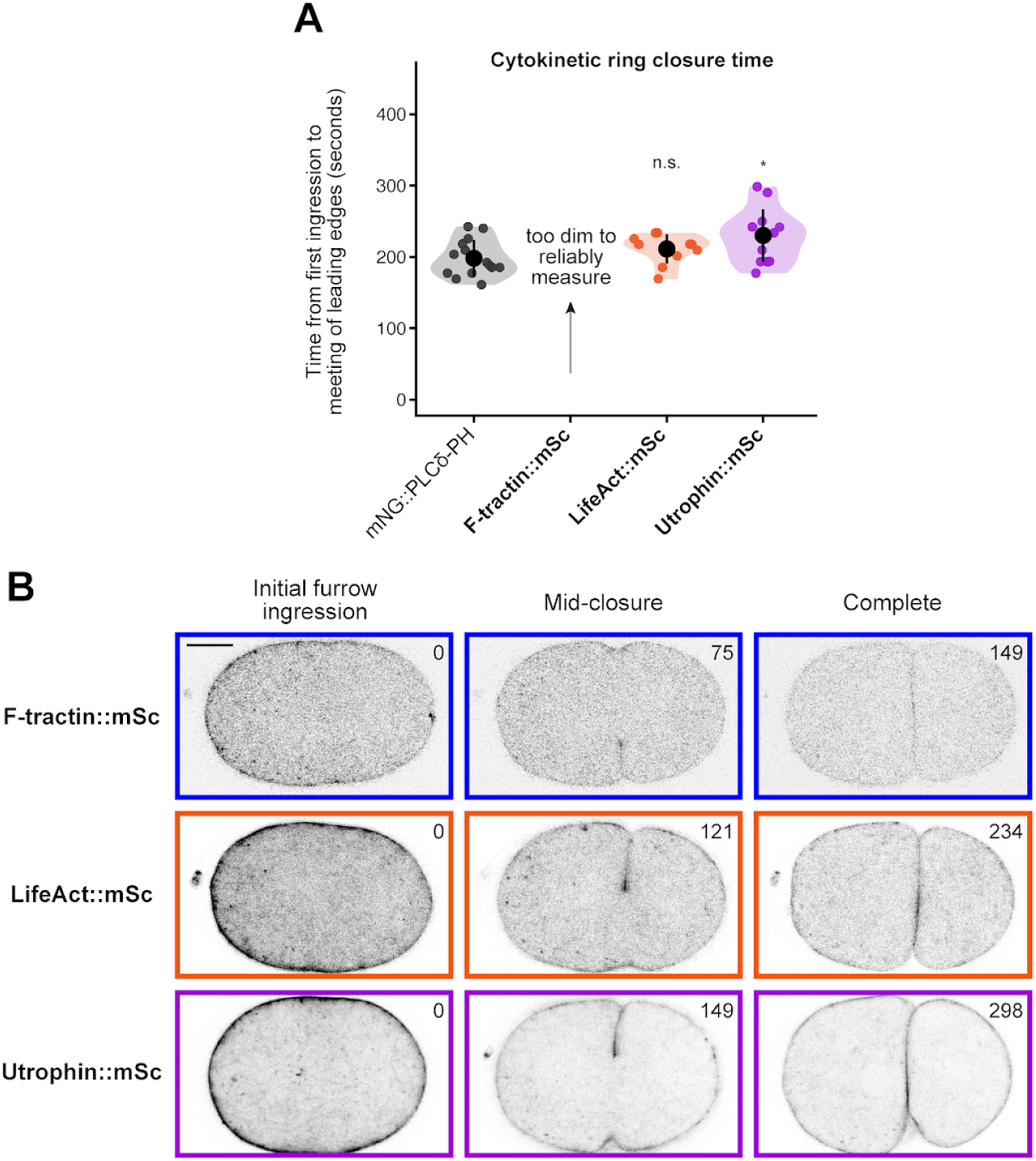
Cytokinetic ring closure in embryos expressing mScarlet-I-tagged F-actin probes. (A) Quantification of cytokinetic ring closure time, measured in seconds from initial furrow ingression to the meeting of the leading edges, in embryos expressing the three mScarlet-I-tagged F-actin probes or the *mex-5p*::mNG::PLCδ-PH control. Statistical significance was determined using one-way ANOVA followed by Dunnett’s post-hoc test, comparing each strain to the *mex-5p*::mNG::PLCδ-PH control. (B) Representative micrographs of embryos expressing the three mScarlet-I-tagged F-actin probes at initial furrow ingression, mid-closure, and completion of cytokinetic ring closure. Fluorescence signals are displayed using an inverted lookup table. Time in seconds is indicated in each image.

**Supplemental Figure S4.**
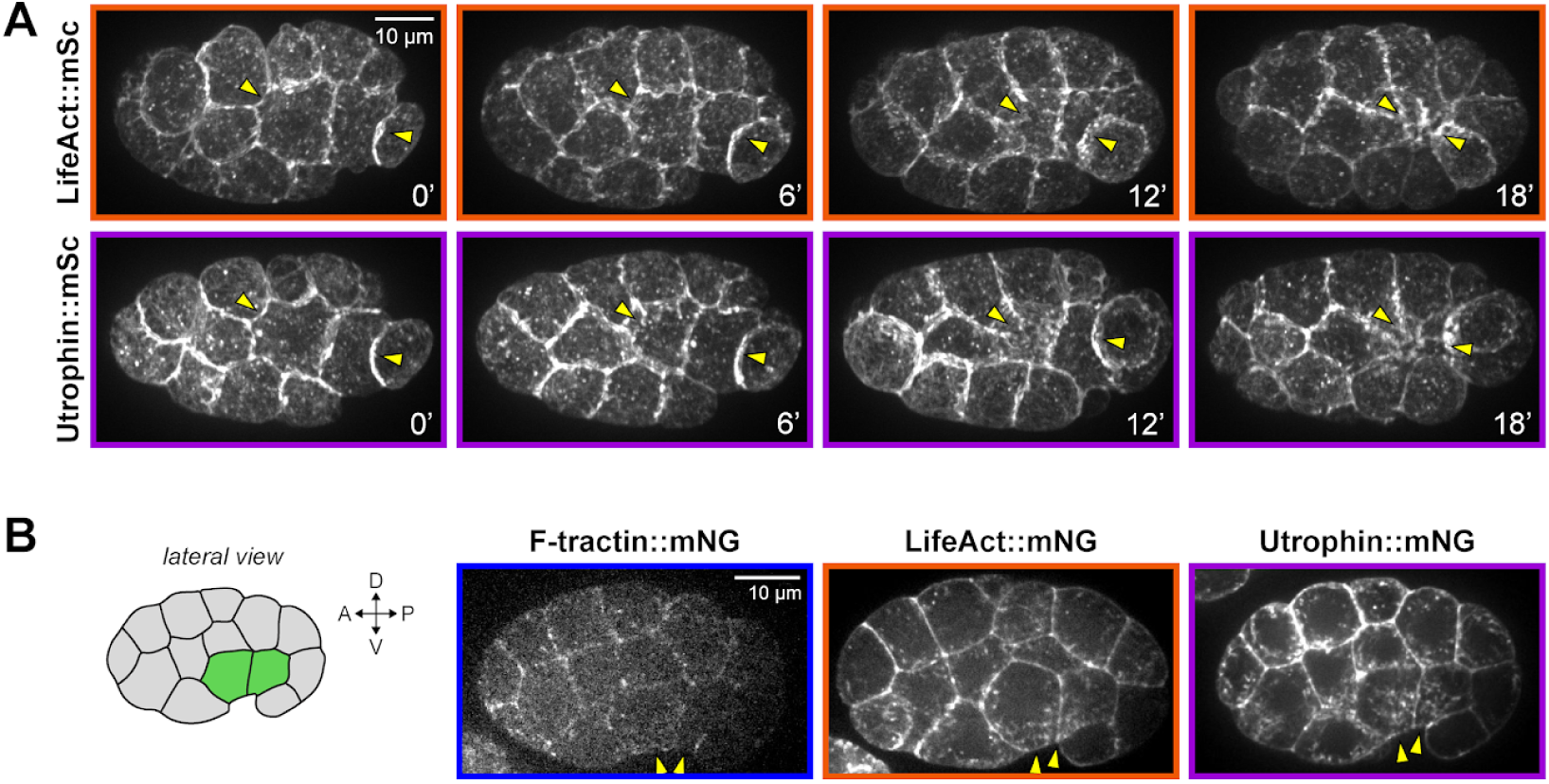
Imaging mScarlet-I- and mNeonGreen-tagged F-actin probes during apical constriction. (A) Representative images of embryos expressing LifeAct::mScarlet-I or Utrophin::mScarlet-I during apical constriction, shown at the same time points as in Figure 6. F-tractin::mScarlet-I fluorescence was too dim to obtain useful time-lapse movies. (B) Representative lateral views of embryos expressing the three mNeonGreen-tagged F-actin probes.

